# Manifold learning uncovers nonlinear interactions between the adolescent brain and environment that predict emotional and behavioral problems

**DOI:** 10.1101/2024.02.29.582854

**Authors:** Erica L. Busch, May I. Conley, Arielle Baskin-Sommers

## Abstract

**Background:** To progress adolescent mental health research beyond our present achievements – a complex account of brain and environmental risk factors without understanding neurobiological embedding in the environment – we need methods to unveil relationships between the developing brain and real-world environmental experiences.

**Methods:** We investigated associations among brain function, environments, and emotional and behavioral problems using participants from the Adolescent Brain and Cognitive Development Study (N=2,401 female). We applied manifold learning, a promising technique for uncovering latent structure from high-dimensional biomedical data like functional magnetic resonance imaging (fMRI). Specifically, we developed *exogenous* PHATE (E-PHATE) to model brain–environment interactions. We used E-PHATE embeddings of participants’ brain activation during emotional and cognitive processing to predict individual differences in cognition and emotional and behavioral problems, both cross-sectionally and longitudinally.

**Results:** E-PHATE embeddings of participants’ brain activation and environments at baseline show moderate-to-large associations with total, externalizing, and internalizing problems at baseline, across several subcortical regions and large-scale cortical networks, relative to the zero-to-small effects achieved by voxel or PHATE methods. E-PHATE embeddings of the brain and environment at baseline also relate to emotional and behavioral problems two years later. These longitudinal predictions show a consistent, moderate effect in the frontoparietal and attention networks.

**Conclusions:** Adolescent brain’s embedding in the environment yields enriched insight into emotional and behavioral problems. Using E-PHATE, we demonstrate how the harmonization of cutting-edge computational methods with longstanding developmental theories advances detection and prediction of adolescent emotional and behavioral problems.

## Introduction

Nearly 75% of all mental health disorders onset during adolescence, with half of all mental health disorders occurring by age 14 (1). Adolescents who experience mental health problems are at heightened risk for lifelong challenges including lower educational attainment, increased legal system involvement, and chronic physical and mental health problems (2). Given the impacts of mental health problems on individuals and society, developmental scientists have long grappled with understanding the emergence of emotional and behavioral problems in youth.

An extensive body of research identifies factors related to emotional and behavioral problems that span neurobiological and environmental factors(3,4). Much of this work has been siloed into work specifying the neurobiology or the environmental factors related mental health problems in adolescence. Neurobiological theories of emotional and behavior problems have emphasized that three key brain regions are especially sensitive during adolescent development: prefrontal cortex (PFC), amygdala, and hippocampus (5). These brain regions support self-regulation and affective processing (6,7) and differences in their functional activation have been related various aspects of emotional and behavioral problems (5). Other research identifies environmental exposures that increase risk for the development of emotional and behavioral problems (8). Meta-analyses report medium to large effects between adversity in adolescents’ families (e.g., conflict, caregiver nonacceptance) and neighborhoods (e.g., experiencing violence or disadvantage) and emotional and behavioral problems (9–11).

Some of the environmental risk factors (e.g., parenting styles, community disadvantage) have been related to the function of mental health-related neural regions (12,13). For instance, a recent study found that the interaction of neighborhood adversity and lower executive network activation during an emotional working memory task was related to higher externalizing problems in adolescents(14). Additional work found that the interaction between neighborhood adversity and decreased amygdala activation during an emotional introspection task was related to higher externalizing problems in a sample of Mexican-origin adolescents(15). Furthermore, another study found that neighborhood and family adversity interacts with PFC functional connectivity to predict internalizing symptoms(16). Across these studies, we can stitch together a model of emotional and behavioral problems that includes interactions between experiences in adolescent’s environments and brain function in regions involved in emotion processing and cognition. Yet, most previous research has modeled this interaction as a linear combination between univariate measures of brain and environment—in other words, considering a single measurement of environment and a univariate signal of brain activation. Some recent work has used multivariate approaches, such as canonical correlation analysis, partial least squares regression, or PCA ridge regression, to maximize brain-behavior associations(17–21). However, these approaches focus on learning components of a multivariate neural representation that are maximally predictive of a target behavior, which may not necessarily reflect the intrinsic geometry of brain activation itself that could arise through unsupervised methods. Further, they frequently fail to account for the multivariate measures of the environment in conjunction with the neural data.

Studying the nonlinear, multidimensional interplay between adolescent brains and environments risk requires computational methods to combine and unveil structure in high-dimensional, multi-modal data. Manifold learning is increasingly popular for highlighting complex latent structure in high-dimensional biological data(22). Manifold learning is an unsupervised, data-driven approach where the dimensionality reduction step is not optimized to maximize a prediction, but rather discovers a manifold of given data. Downstream, one can use manifold embeddings to test associations between the manifold and other information and gauge the quality or type of information represented within the manifold. The algorithm PHATE was specifically designed for high-dimensional, noisy biomedical data and has been applied to uncover local and global latent structure in functional magnetic resonance imaging (fMRI) data(23–25). Prior research has shown that combining PHATE with additional data (e.g., temporal dynamics of brain responses) enhances the relevance of embeddings for understanding complex cognitive processing (i.e., during movie viewing)(26). However, standard PHATE implementations cannot account for interactions of additional variables outside of the high-dimensional input data (e.g., brain data).

Here, using the Adolescent Brain Cognitive Development^SM^ Study (ABCD Study®) baseline sample and two-year data, we investigated the interplay of environment and brain function on emotional and behavioral problems. We tested 1) whether PHATE can be used to enhance the behavioral relevance of task-based, developmental fMRI data and 2) whether an updated version of PHATE can be combined with environmental data to discover latent geometric structure connecting adolescents’ brains, environments, and emotional and behavioral problems. First, as a proof of concept, we showed that PHATE embeddings of brain activation during cognitive and emotion processing(27) were strongly associated with individual differences in working memory performance in 9—10 year-olds. Next, we combined the PHATE brain activation manifold with measurements of adolescents’ environments into a *multi-view* manifold. Multi-view approaches combine different measurements collected from the same samples into a single representation to be embedded in lower dimensions. For instance, temporal PHATE (T-PHATE) is a recently-introduced multi-view algorithm combining two signals *endogenous* to brain data (i.e., calculated directly from the fMRI measurements)(26). In the present study, we introduce *exogenous* PHATE (E-PHATE), which combines participants’ PHATE brain activation manifold with data about the same participants collected *externally* (i.e., family and neighborhood adversity). We hypothesized that the development of emotional and behavioral problems would relate to a nonlinear interaction between the adolescent brain and their environments. E-PHATE embeddings showed a stronger relationship with emotional and behavioral problems both cross-sectionally and longitudinally than either the PHATE or original voxel-wise data(3,4,28,29). E-PHATE sheds light on the neural-environment interplay and improves the detection and prediction of emotional and behavioral problems in adolescents.

## Methods and materials

### Participants

Participants were adolescents included in ABCD Data Release 4.0 (DOI:10.15154/1523041). Neuroimaging and environment data were from the baseline assessment (ages 9–10). Emotional and behavioral data were from the baseline and the 2-year follow up (ages 11-12). Participants were excluded for missing fMRI, environmental, or mental health measures, resulting in 4,732 participants included in baseline analyses and 2,371 participants included in longitudinal analyses (Table 1 for sample demographics; Supplemental Methods S1, Table S3 & S4, and Figure S1 for participant selection).

**Table 1.** Demographics for participants included in the baseline (ages 9-10) and 2-year follow-up (ages 11-12) analyses.

| **Timepoint** | **Baseline** | **2-year follow-up** |
| --- | --- | --- |
| **n (percent)** | 4,372 | 2,371 |
| **Female** | 2,401 (51) | 1,338 (49) |
| **Race/Ethnicity** |  |  |
| Hispanic or Latinx | 883 (18.7) | 457 (18.6) |
| Black | 518 (10.9) | 232 (9.1) |
| White | 2710 (57.2) | 1,545 (60.7) |
| Asian | 107 (2.26) | 51 (2.0) |
| Other | 513 (10.8) | 259 (10.1) |
| **Caregiver education (%)** |  |  |
| < HS Diploma | 207 (4.4) | 98 (3.9) |
| HS Diploma / GED | 426 (9.0) | 214 (8.4) |
| Some College | 1,299 (27.5) | 704 (27.7) |
| Bachelor | 1,463 (30.9) | 818 (32.2) |
| Post Graduate Degree | 1334 (28.2) | 708 (27.8) |
| Unknown | 3 (0.06) | 2 (0.08) |
| **Family income (%)** |  |  |
| < \$50K | 1,047 (22.1) | 538 (21.1) |
| > \$50K - < \$100K | 1,298 (27.4) | 752 (29.6) |
| > \$100K | 2,053 (43.4) | 1088 (42.8) |
| Unknown | 334 (7.1) | 166 (6.5) |

### Environment measures

Measures of the environment were selected to characterize youth’s family and neighborhood environments at the baseline timepoint. Family environment was measured using participants’ perception of family threat (anger and conflict expressed among family members) and of family support (caregiver acceptance). Neighborhood environment was measured using participant and caregiver assessment of perceived safety/crime, and the Area Deprivation Index (ADI), a composite index of neighborhood socioeconomic disadvantage (Supplemental Methods S2).

### Emotional and behavioral measures

Measures of emotional and behavioral problems were assessed using *t*-scores from the baseline and 2-year follow-up data from the Achenbach System of Empirically Based Assessment Child Behavior Checklist (CBCL), which is a 119 item parent/caregiver-report survey of adolescent emotional and behavioral problems validated for use in children ages 6–18(30). Primary analyses examined total problems and externalizing and internalizing broad-band scales. Supplemental analyses examined anxious/depression, withdrawn/depression, somatization, aggression, and rule-breaking behavior syndrome scales.

### Neuroimaging task data

The in-scanner emotional n-back (EN-back) task was designed to engage emotion and memory processing(*27,31*). During each fMRI run, participants performed four 0-back (low memory load) and four 2-back (high memory load) blocks with happy, fear, or neutral face or place stimuli. EN-back performance was measured with sensitivity, calculated as d’ = z(hits) - z(false alarms) and adjusted for extreme values using the Hautus adjustment(32,33). Task information is detailed further in Supplemental Methods S3.

fMRI data were preprocessed by the ABCD Study Data Analysis, Informatics, and Resource Center (DAIRC)(34). EN-back fMRI activation was estimated for each participant using general linear models. Following prior studies, cognitive processing activation was measured as the contrast of 2-back and 0-back blocks, and emotion processing activation was measured as the contrast of emotional and neutral face blocks(14,35). Further information about acquisition and processing are in Supplemental Methods S4.

For each contrast, we analyzed beta weights from cortical networks (defined using the Yeo 7 network solution(36) and the Shaefer 400 parcellation(37)) and subcortical regions (defined using the Scale I Tian subcortical parcellation(38)). We then extracted one beta weight per voxel within each region or network and then vectorized the multi-voxel beta weights, resulting in one vector of voxel-wise beta weights for each region, participant, and contrast. All participants’ vectors are stacked into a single matrix for each region and contrast, and henceforth we refer to these matrices as “voxel-wise data.”

### PHATE manifold learning

We tested whether manifold learning uncovers behaviorally relevant brain activation during cognitive and emotional processing, which could be used to improve prediction of emotional and behavioral problems. First, we applied the PHATE algorithm. PHATE embeddings denoise and highlight local and global nonlinear structure among data points in a low-dimensional representation. Prior work has shown that PHATE embeddings of fMRI data improve sensitivity of the data for predicting features such as functional brain maturity(25) and visual category information(24,26). Using PHATE, we embedded the voxel-wise data for each region and contrast into lower dimensions to test whether the PHATE embeddings uncover individual differences in brain function related to cognition more strongly than the voxel-wise data.

### Exogenous PHATE (E-PHATE) manifold learning

We designed *exogenous* PHATE (E-PHATE) to model this interplay as a low-dimensional manifold. We applied a dual-diffusion(26) approach to combine *exogenous* information about adolescents’ environments with their brain activation manifold. The E-PHATE procedure starts by calculating a PHATE diffusion matrix over the voxel-wise data, which can be considered the affinity among participants’ voxel-wise activation vectors. E-PHATE also calculates a second affinity matrix over those participants’ scores on additional, *exogenous* variables (i.e., family conflict, caregiver acceptance, youth and caregiver perceived neighborhood crime/safety, and neighborhood disadvantage), and these views are combined using dual-diffusion to calculate the E-PHATE diffusion matrix. This matrix is then embedded into *D* dimensions using multidimensional scaling (where D ∈ ℕ; D ∈ {2, 3} for visualization) (Figure 1; Supplemental Methods S5 for the algorithm).

**Figure 1.**
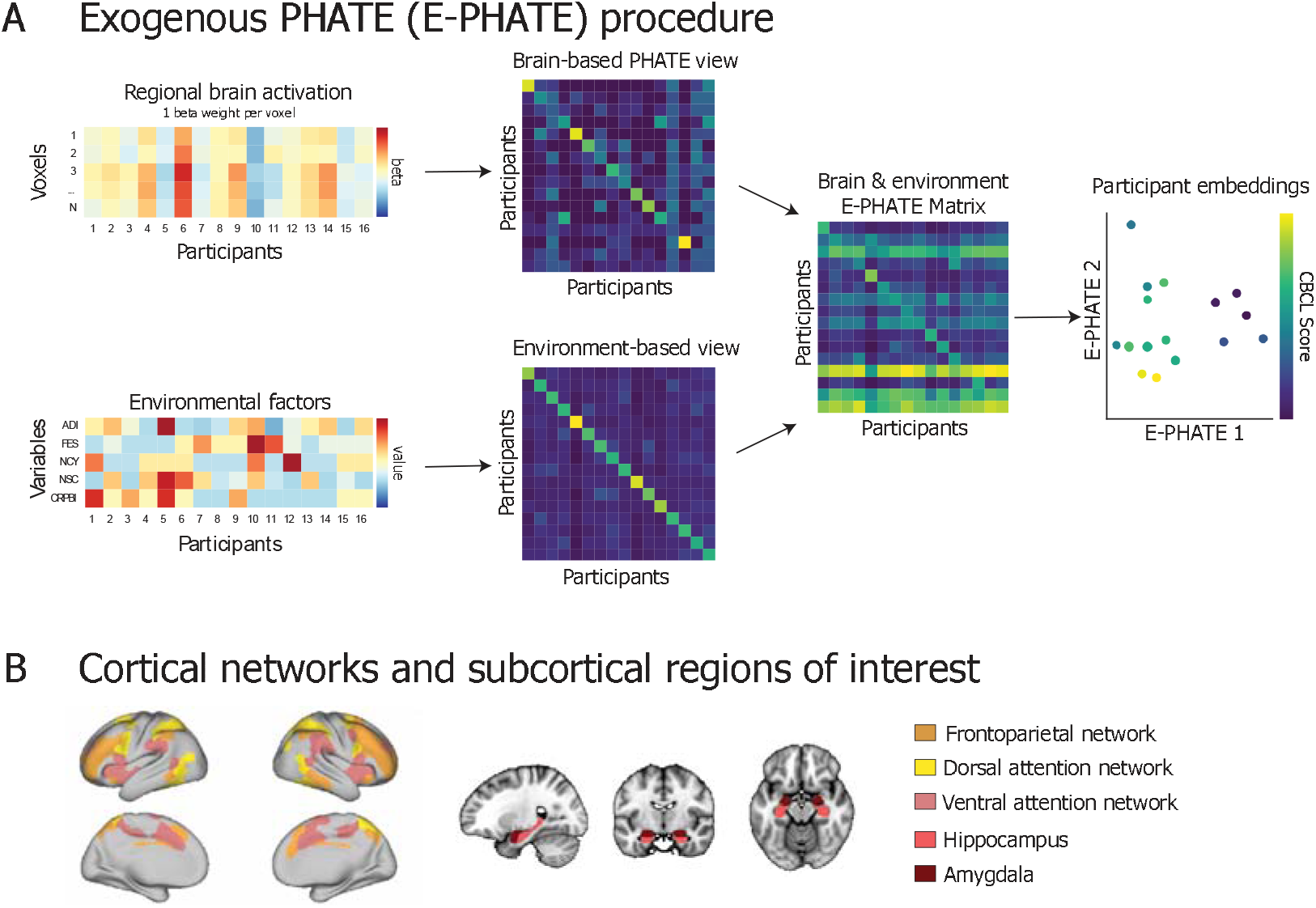
Exogenous PHATE (E-PHATE) procedure. **A:** E-PHATE models the interactions between brain activation and exogenous information about participants using multi-view manifold learning. In this schematic, the first view of E-PHATE takes as inputs a vector of voxel-wise beta values for each participant and computes a PHATE-based affinity matrix between participants’ brain activations. The second view takes a vector of environment scores for each of those participants and builds an affinity matrix across those scores. Both matrices row-normalized to become transition probability matrices. These two views are combined into the E-PHATE diffusion matrix, which now captures both brain and environmental relations among participants. The E-PHATE matrix is then embedded using metric multidimensional scaling (m-MDS). Two dimensions and a subset of participants are shown for visualization; 20 dimensions were used for the main analysis. Participants’ coordinates in E-PHATE dimensions visually reflect individual differences along emotional and behavioral problems (e.g., externalizing problem scores). ADI = area deprivation index, a composite index of neighborhood disadvantage; FES = family conflict; NCY = youth perceived neighborhood safety/crime; NSC = caregiver perceived neighborhood safety/crime; CRPBI = caregiver acceptance **B:** Analyses are presented using beta values extracted for voxels in the bilateral amygdala and hippocampus and surface vertices in three cortical networks: frontoparietal, dorsal and ventral attention networks.

### Benchmarking manifold learning methods

In main analyses, we benchmarked the relevance of E-PHATE and PHATE embeddings against their corresponding (i.e., within region and contrast) voxel-wise data. In supplemental analyses, we benchmarked E-PHATE with principal components analysis (PCA; a linear dimensionality reduction method) and universal manifold approximation and projection (UMAP(39); a common nonlinear embedding method) (Supplemental Methods S7). We also benchmarked E-PHATE, as described above, with variants to test the impact of specific environment variables and algorithmic choices (Supplemental Methods S8—S9). As in prior works, all analyzed embeddings are 20 dimensional for consistency across methods and regions(23,26). Voxel-wise data dimensionality can be found in Supplemental Table 2.

### Prediction of emotional and behavior problems and task performance from brain data

Cross-sectional analyses used participants’ brain data at baseline to predict behavioral scores at baseline (N=4,732). Longitudinal analyses used brain data at baseline to predict scores at the 2-year follow up, controlling for the participant’s score at baseline (N=2,371). For each cross-validation fold, multiple linear regressions were trained on 95% of participants’ brain data (either voxel-wise or embedding) to predict participants’ EN-back task or CBCL scores(20,21,26). Regressions were then applied to predict scores from the brain data (i.e., voxel-wise or embedding) of held-out participants and scored as the partial Spearman’s correlation (p) between predicted and true scores for each fold of held-out participants. p values were then averaged across folds. In our analyses, we compared the partial Spearman’s p of multiple linear regression models trained on voxel-wise data, PHATE, and E-PHATE embeddings. Covariates included scanner serial number for cross-sectional and longitudinal analyses and participants’ baseline scores for the behavioral measure being predicted in longitudinal analyses. Performance of regression models trained on different data representations (e.g., voxel-wise vs. E-PHATE) was compared across representations using pairwise permutation tests (10,000 iterations) and Bonferroni correction; comparisons across representations refer these p-values. Further details about regression models and statistical testing are included in Supplemental Methods S6 and S10, respectively. Analysis code and software for E-PHATE can be found <u>here</u>. E-PHATE will be released as a python package available with pip upon publication, and all code will be made publicly available on Github.

## Results

### PHATE strengthened associations between brain activation and EN-back performance

As a proof-of-concept to confirm if manifold learning could improve representation of cognitively-relevant brain activity, we first tested the association between standard PHATE embeddings with participants’ EN-back task scores. Voxel-wise 2-back vs. 0-back activation showed moderate associations with EN-back performance (p < 0.20) in all regions (Table 2). PHATE embeddings of 2-back vs. 0-back activation were significantly related to EN-back task performance in all regions (Figure 2, Table 2; Supplemental Data 2) with a large effect size (p > 0.20). The greatest effects were observed in the in the frontoparietal and attention networks (p > 0.52), which were over double the effect sizes (p) for regression models voxel-wise data. Given previous research linking frontoparietal and attention networks with higher order cognitive abilities and working memory(35,40,41), these results demonstrate that PHATE optimized the sensitivity of the fMRI data for detecting brain activation related to cognitive performance. Consistent with prior research(35), none of the voxel-wise data for the emotion vs. neutral face contrast significantly related to EN-back performance. In contrast, PHATE embeddings of emotion vs. neutral activation showed moderate associations with EN-back performance (p *>* 0.14 in frontoparietal and attention networks). Therefore, the PHATE embedding enhanced access to brain activation related to emotion processing during the EN-back task.

**Figure 2:**
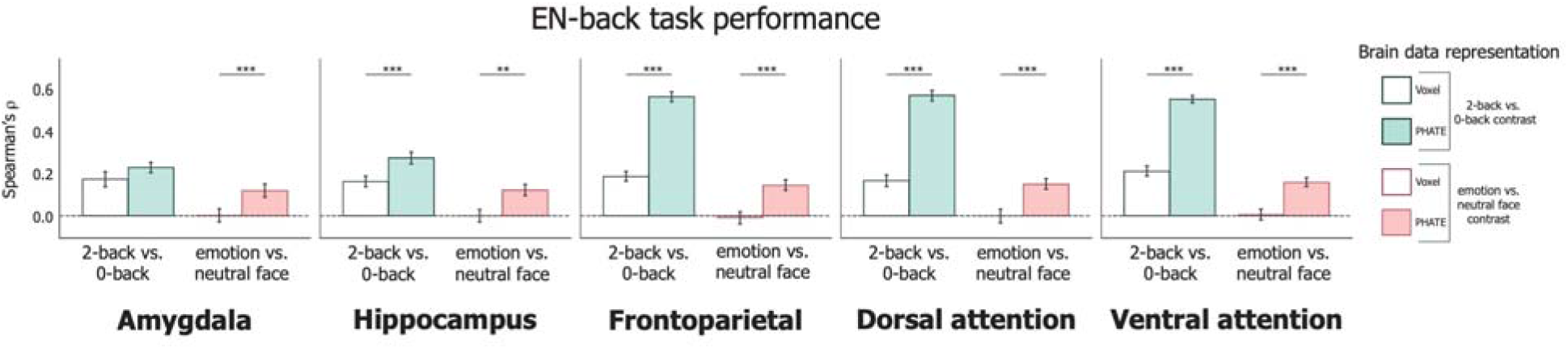
Associations of brain data with EN-back task performance. Multiple linear regression was used to measure the association between brain data representations and EN-back task performance and scored as the partial Spearman correlation (p) between true and regression-predicted performance for held-out participants (20 cross-validation folds). Bars represent the average p across cross-validation test folds. Pairwise permutation tests were used to compute p-values for the difference in p for each cross-validation fold across representations of brain data (i.e., voxel-resolution data in white and corresponding PHATE embedding filled in). Left bar set in each graph represents the 2-back vs. 0-back contrast; Right bar set in each graph represents the emotion vs. neutral contrast. A=Total problems; B=Externalizing problems; C=Internalizing problems. Error bars represent the 95% confidence interval of the mean p across 20 cross-validation folds. ∼ p < 0.1, * p < 0.05, ** p < 0.01, *** p < 0.001

**Table 2.**
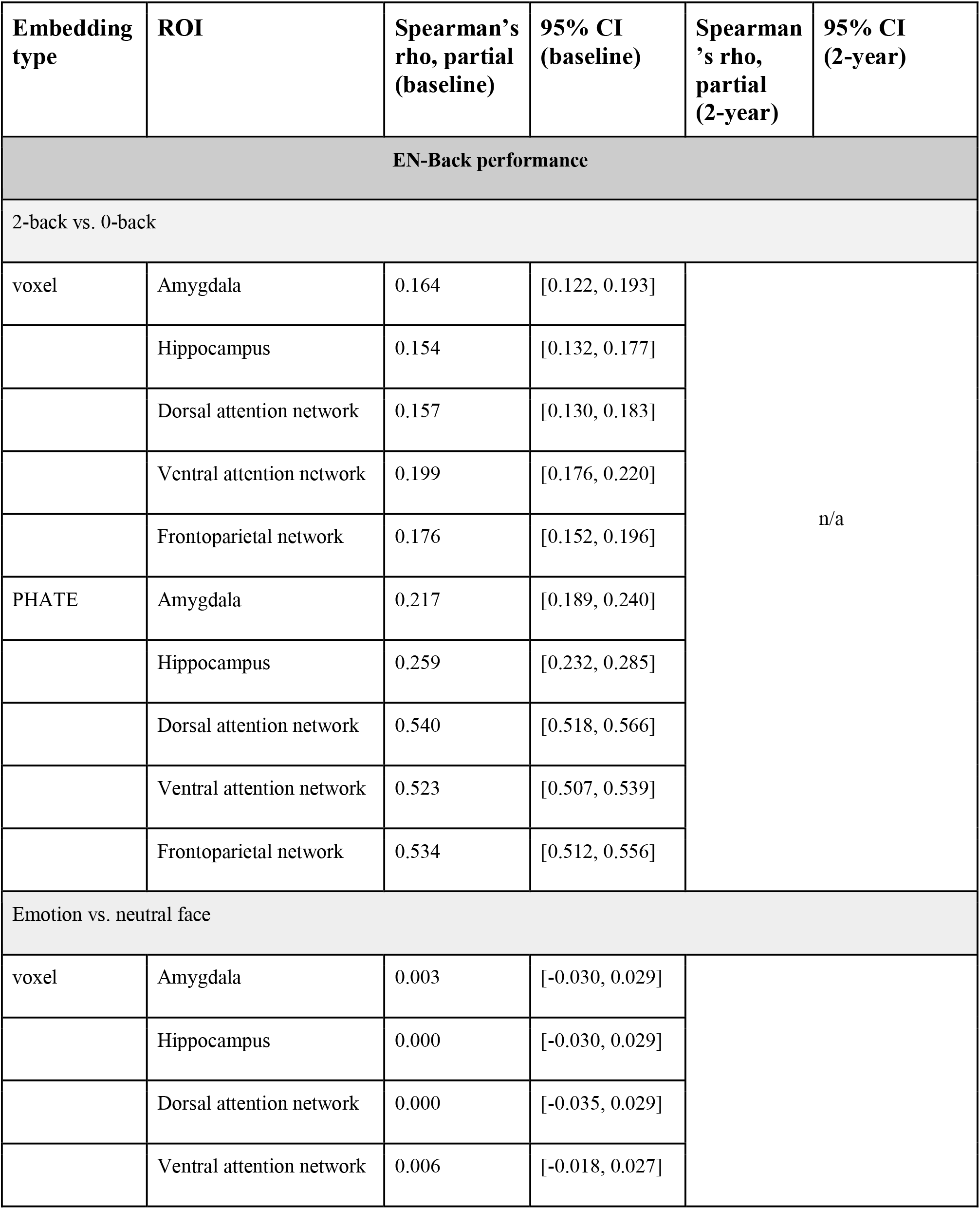
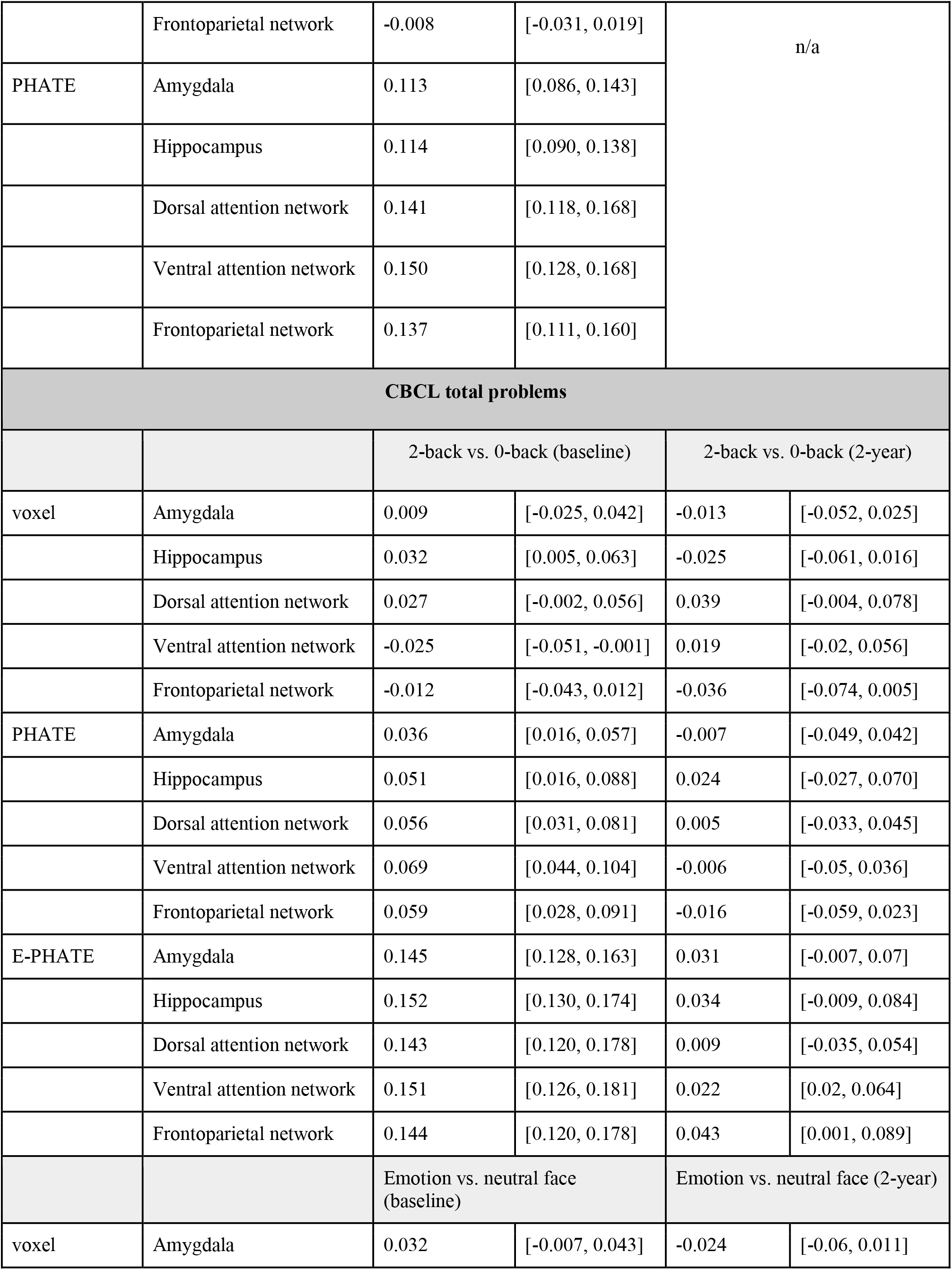
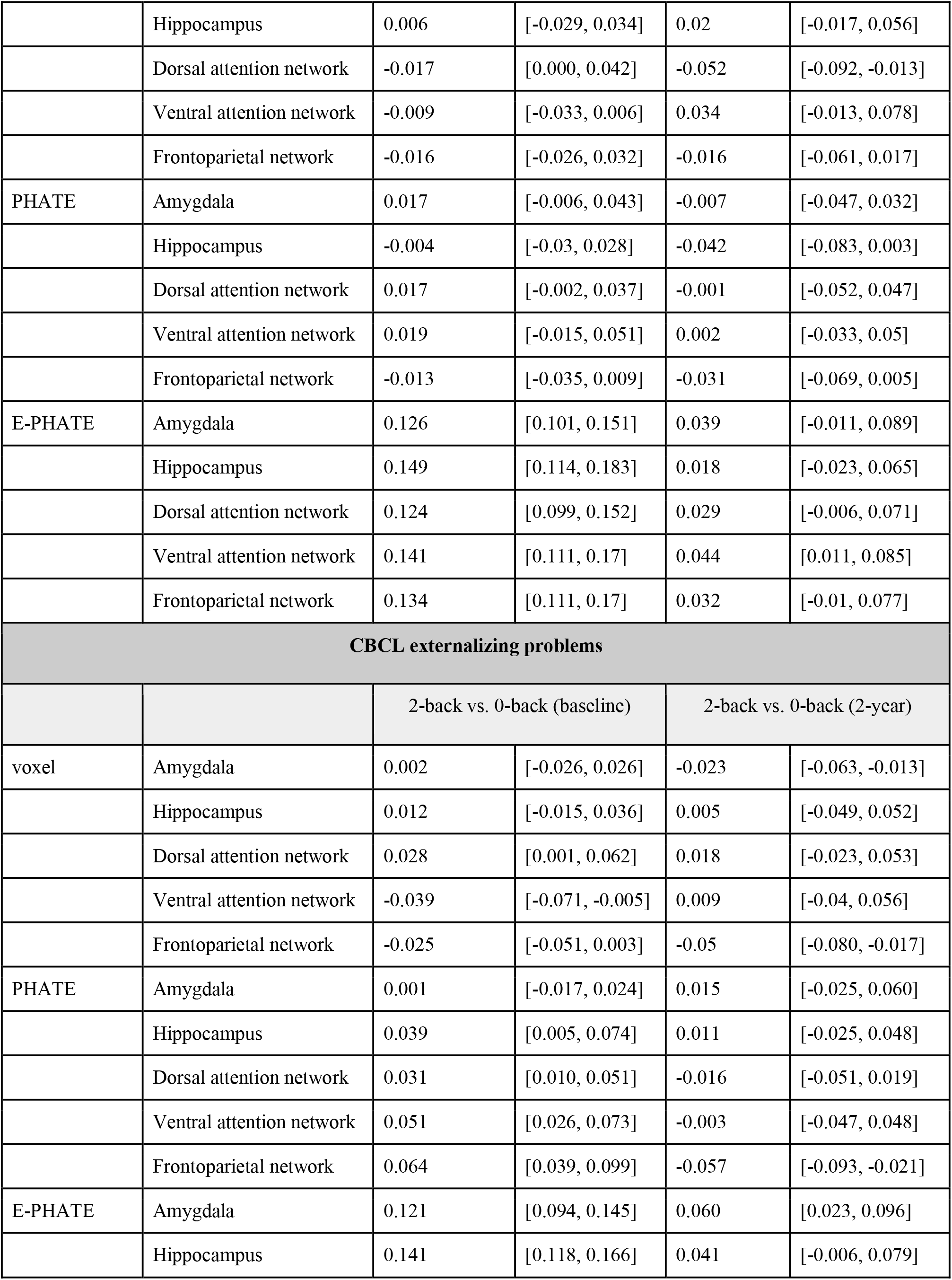
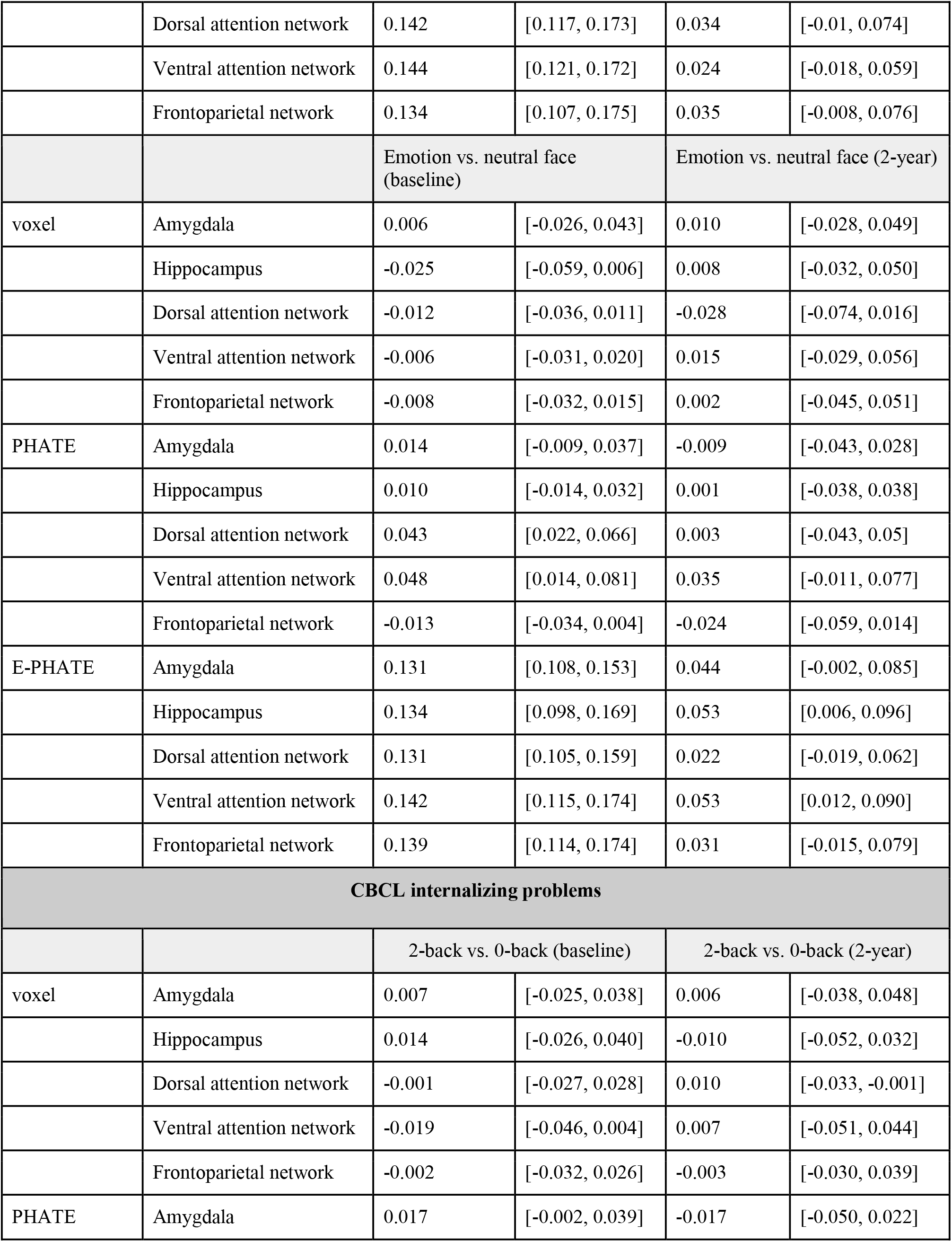
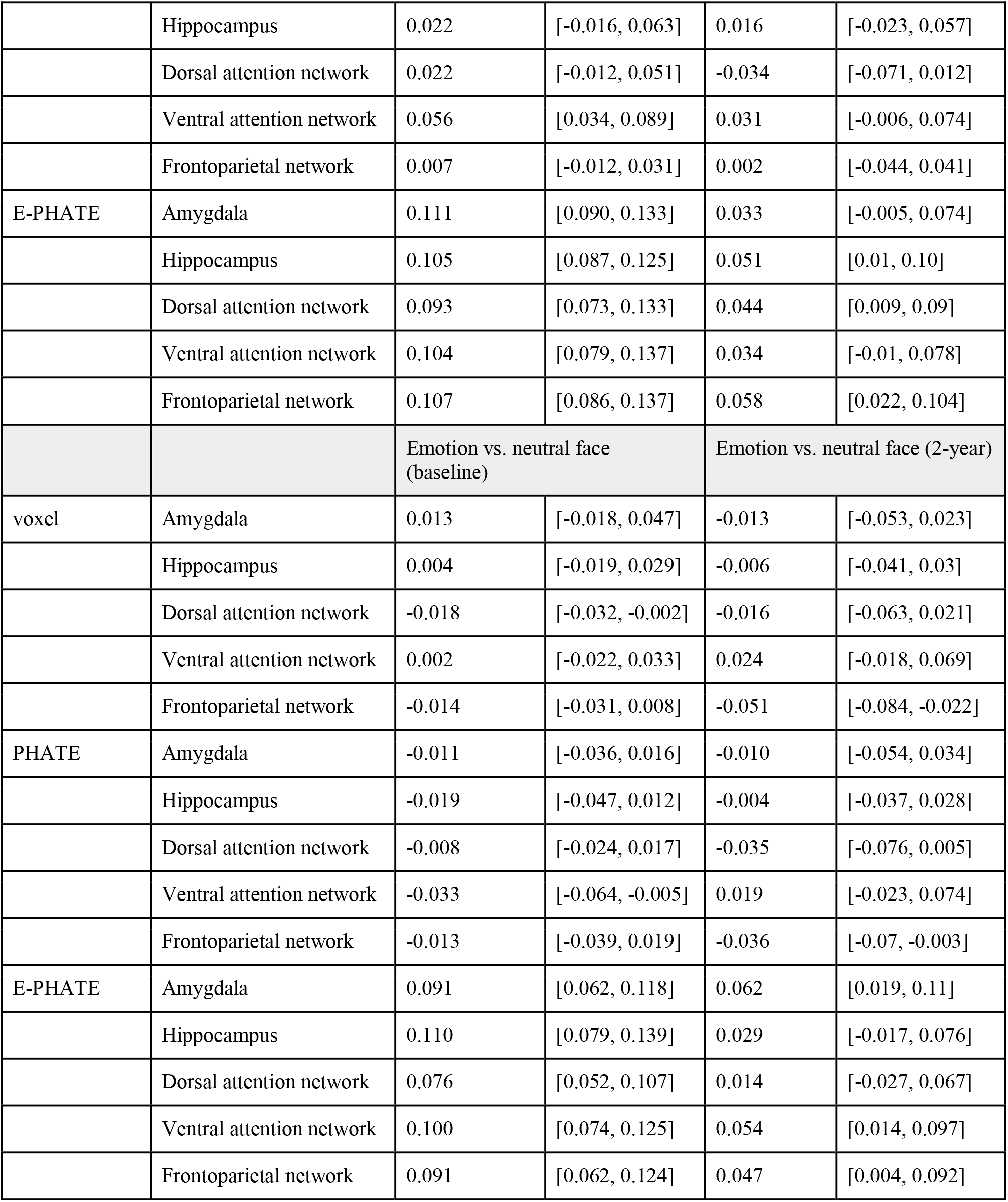
Results showing associations between brain activation embeddings and task performance and emotional and behavioral problems.

| Embedding type | ROI | Spearman’s rho, partial (baseline) | 95% CI (baseline) | Spearman’s rho, partial (2-year) | 95% CI (2-year) |
| --- | --- | --- | --- | --- | --- |
| EN-Back performance |  |  |  |  |  |
| 2-back vs. 0-back |  |  |  |  |  |
| voxel | Amygdala | 0.164 | [0.122, 0.193] | n/a |  |
|  | Hippocampus | 0.154 | [0.132, 0.177] |  |  |
|  | Dorsal attention network | 0.157 | [0.130, 0.183] |  |  |
|  | Ventral attention network | 0.199 | [0.176, 0.220] |  |  |
|  | Frontoparietal network | 0.176 | [0.152, 0.196] |  |  |
| PHATE | Amygdala | 0.217 | [0.189, 0.240] |  |  |
|  | Hippocampus | 0.259 | [0.232, 0.285] |  |  |
|  | Dorsal attention network | 0.540 | [0.518, 0.566] |  |  |
|  | Ventral attention network | 0.523 | [0.507, 0.539] |  |  |
|  | Frontoparietal network | 0.534 | [0.512, 0.556] |  |  |
| Emotion vs. neutral face |  |  |  |  |  |
| voxel | Amygdala | 0.003 | [-0.030, 0.029] |  |  |
|  | Hippocampus | 0.000 | [-0.030, 0.029] |  |  |
|  | Dorsal attention network | 0.000 | [-0.035, 0.029] |  |  |
|  | Ventral attention network | 0.006 | [-0.018, 0.027] |  |  |
|  | Frontoparietal network | -0.008 | [-0.031, 0.019] | n/a |  |
| PHATE | Amygdala | 0.113 | [0.086, 0.143] |  |  |
|  | Hippocampus | 0.114 | [0.090, 0.138] |  |  |
|  | Dorsal attention network | 0.141 | [0.118, 0.168] |  |  |
|  | Ventral attention network | 0.150 | [0.128, 0.168] |  |  |
|  | Frontoparietal network | 0.137 | [0.111, 0.160] |  |  |
| CBCL total problems |  |  |  |  |  |
|  |  | 2-back vs. 0-back (baseline) |  | 2-back vs. 0-back (2-year) |  |
| voxel | Amygdala | 0.009 | [-0.025, 0.042] | -0.013 | [-0.052, 0.025] |
|  | Hippocampus | 0.032 | [0.005, 0.063] | -0.025 | [-0.061, 0.016] |
|  | Dorsal attention network | 0.027 | [-0.002, 0.056] | 0.039 | [-0.004, 0.078] |
|  | Ventral attention network | -0.025 | [-0.051, -0.001] | 0.019 | [-0.02, 0.056] |
|  | Frontoparietal network | -0.012 | [-0.043, 0.012] | -0.036 | [-0.074, 0.005] |
| PHATE | Amygdala | 0.036 | [0.016, 0.057] | -0.007 | [-0.049, 0.042] |
|  | Hippocampus | 0.051 | [0.016, 0.088] | 0.024 | [-0.027, 0.070] |
|  | Dorsal attention network | 0.056 | [0.031, 0.081] | 0.005 | [-0.033, 0.045] |
|  | Ventral attention network | 0.069 | [0.044, 0.104] | -0.006 | [-0.05, 0.036] |
|  | Frontoparietal network | 0.059 | [0.028, 0.091] | -0.016 | [-0.059, 0.023] |
| E-PHATE | Amygdala | 0.145 | [0.128, 0.163] | 0.031 | [-0.007, 0.07] |
|  | Hippocampus | 0.152 | [0.130, 0.174] | 0.034 | [-0.009, 0.084] |
|  | Dorsal attention network | 0.143 | [0.120, 0.178] | 0.009 | [-0.035, 0.054] |
|  | Ventral attention network | 0.151 | [0.126, 0.181] | 0.022 | [0.02, 0.064] |
|  | Frontoparietal network | 0.144 | [0.120, 0.178] | 0.043 | [0.001, 0.089] |
|  |  | Emotion vs. neutral face (baseline) |  | Emotion vs. neutral face (2-year) |  |
| voxel | Amygdala | 0.032 | [-0.007, 0.043] | -0.024 | [-0.06, 0.011] |
|  | Hippocampus | 0.006 | [-0.029, 0.034] | 0.02 | [-0.017, 0.056] |
|  | Dorsal attention network | -0.017 | [0.000, 0.042] | -0.052 | [-0.092, -0.013] |
|  | Ventral attention network | -0.009 | [-0.033, 0.006] | 0.034 | [-0.013, 0.078] |
|  | Frontoparietal network | -0.016 | [-0.026, 0.032] | -0.016 | [-0.061, 0.017] |
| PHATE | Amygdala | 0.017 | [-0.006, 0.043] | -0.007 | [-0.047, 0.032] |
|  | Hippocampus | -0.004 | [-0.03, 0.028] | -0.042 | [-0.083, 0.003] |
|  | Dorsal attention network | 0.017 | [-0.002, 0.037] | -0.001 | [-0.052, 0.047] |
|  | Ventral attention network | 0.019 | [-0.015, 0.051] | 0.002 | [-0.033, 0.05] |
|  | Frontoparietal network | -0.013 | [-0.035, 0.009] | -0.031 | [-0.069, 0.005] |
| E-PHATE | Amygdala | 0.126 | [0.101, 0.151] | 0.039 | [-0.011, 0.089] |
|  | Hippocampus | 0.149 | [0.114, 0.183] | 0.018 | [-0.023, 0.065] |
|  | Dorsal attention network | 0.124 | [0.099, 0.152] | 0.029 | [-0.006, 0.071] |
|  | Ventral attention network | 0.141 | [0.111, 0.17] | 0.044 | [0.011, 0.085] |
|  | Frontoparietal network | 0.134 | [0.111, 0.17] | 0.032 | [-0.01, 0.077] |
| **CBCL externalizing problems** |  |  |  |  |  |
|  |  | 2-back vs. 0-back (baseline) |  | 2-back vs. 0-back (2-year) |  |
| voxel | Amygdala | 0.002 | [-0.026, 0.026] | -0.023 | [-0.063, -0.013] |
|  | Hippocampus | 0.012 | [-0.015, 0.036] | 0.005 | [-0.049, 0.052] |
|  | Dorsal attention network | 0.028 | [0.001, 0.062] | 0.018 | [-0.023, 0.053] |
|  | Ventral attention network | -0.039 | [-0.071, -0.005] | 0.009 | [-0.04, 0.056] |
|  | Frontoparietal network | -0.025 | [-0.051, 0.003] | -0.05 | [-0.080, -0.017] |
| PHATE | Amygdala | 0.001 | [-0.017, 0.024] | 0.015 | [-0.025, 0.060] |
|  | Hippocampus | 0.039 | [0.005, 0.074] | 0.011 | [-0.025, 0.048] |
|  | Dorsal attention network | 0.031 | [0.010, 0.051] | -0.016 | [-0.051, 0.019] |
|  | Ventral attention network | 0.051 | [0.026, 0.073] | -0.003 | [-0.047, 0.048] |
|  | Frontoparietal network | 0.064 | [0.039, 0.099] | -0.057 | [-0.093, -0.021] |
| E-PHATE | Amygdala | 0.121 | [0.094, 0.145] | 0.060 | [0.023, 0.096] |
|  | Hippocampus | 0.141 | [0.118, 0.166] | 0.041 | [-0.006, 0.079] |
|  | Dorsal attention network | 0.142 | [0.117, 0.173] | 0.034 | [-0.01, 0.074] |
|  | Ventral attention network | 0.144 | [0.121, 0.172] | 0.024 | [-0.018, 0.059] |
|  | Frontoparietal network | 0.134 | [0.107, 0.175] | 0.035 | [-0.008, 0.076] |
|  |  | Emotion vs. neutral face (baseline) |  | Emotion vs. neutral face (2-year) |  |
| voxel | Amygdala | 0.006 | [-0.026, 0.043] | 0.010 | [-0.028, 0.049] |
|  | Hippocampus | -0.025 | [-0.059, 0.006] | 0.008 | [-0.032, 0.050] |
|  | Dorsal attention network | -0.012 | [-0.036, 0.011] | -0.028 | [-0.074, 0.016] |
|  | Ventral attention network | -0.006 | [-0.031, 0.020] | 0.015 | [-0.029, 0.056] |
|  | Frontoparietal network | -0.008 | [-0.032, 0.015] | 0.002 | [-0.045, 0.051] |
| PHATE | Amygdala | 0.014 | [-0.009, 0.037] | -0.009 | [-0.043, 0.028] |
|  | Hippocampus | 0.010 | [-0.014, 0.032] | 0.001 | [-0.038, 0.038] |
|  | Dorsal attention network | 0.043 | [0.022, 0.066] | 0.003 | [-0.043, 0.05] |
|  | Ventral attention network | 0.048 | [0.014, 0.081] | 0.035 | [-0.011, 0.077] |
|  | Frontoparietal network | -0.013 | [-0.034, 0.004] | -0.024 | [-0.059, 0.014] |
| E-PHATE | Amygdala | 0.131 | [0.108, 0.153] | 0.044 | [-0.002, 0.085] |
|  | Hippocampus | 0.134 | [0.098, 0.169] | 0.053 | [0.006, 0.096] |
|  | Dorsal attention network | 0.131 | [0.105, 0.159] | 0.022 | [-0.019, 0.062] |
|  | Ventral attention network | 0.142 | [0.115, 0.174] | 0.053 | [0.012, 0.090] |
|  | Frontoparietal network | 0.139 | [0.114, 0.174] | 0.031 | [-0.015, 0.079] |
| **CBCL internalizing problems** |  |  |  |  |  |
|  |  | 2-back vs. 0-back (baseline) |  | 2-back vs. 0-back (2-year) |  |
| voxel | Amygdala | 0.007 | [-0.025, 0.038] | 0.006 | [-0.038, 0.048] |
|  | Hippocampus | 0.014 | [-0.026, 0.040] | -0.010 | [-0.052, 0.032] |
|  | Dorsal attention network | -0.001 | [-0.027, 0.028] | 0.010 | [-0.033, -0.001] |
|  | Ventral attention network | -0.019 | [-0.046, 0.004] | 0.007 | [-0.051, 0.044] |
|  | Frontoparietal network | -0.002 | [-0.032, 0.026] | -0.003 | [-0.030, 0.039] |
| PHATE | Amygdala | 0.017 | [-0.002, 0.039] | -0.017 | [-0.050, 0.022] |
|  | Hippocampus | 0.022 | [-0.016, 0.063] | 0.016 | [-0.023, 0.057] |
|  | Dorsal attention network | 0.022 | [-0.012, 0.051] | -0.034 | [-0.071, 0.012] |
|  | Ventral attention network | 0.056 | [0.034, 0.089] | 0.031 | [-0.006, 0.074] |
|  | Frontoparietal network | 0.007 | [-0.012, 0.031] | 0.002 | [-0.044, 0.041] |
| E-PHATE | Amygdala | 0.111 | [0.090, 0.133] | 0.033 | [-0.005, 0.074] |
|  | Hippocampus | 0.105 | [0.087, 0.125] | 0.051 | [0.01, 0.10] |
|  | Dorsal attention network | 0.093 | [0.073, 0.133] | 0.044 | [0.009, 0.09] |
|  | Ventral attention network | 0.104 | [0.079, 0.137] | 0.034 | [-0.01, 0.078] |
|  | Frontoparietal network | 0.107 | [0.086, 0.137] | 0.058 | [0.022, 0.104] |
|  |  | Emotion vs. neutral face (baseline) |  | Emotion vs. neutral face (2-year) |  |
| voxel | Amygdala | 0.013 | [-0.018, 0.047] | -0.013 | [-0.053, 0.023] |
|  | Hippocampus | 0.004 | [-0.019, 0.029] | -0.006 | [-0.041, 0.03] |
|  | Dorsal attention network | -0.018 | [-0.032, -0.002] | -0.016 | [-0.063, 0.021] |
|  | Ventral attention network | 0.002 | [-0.022, 0.033] | 0.024 | [-0.018, 0.069] |
|  | Frontoparietal network | -0.014 | [-0.031, 0.008] | -0.051 | [-0.084, -0.022] |
| PHATE | Amygdala | -0.011 | [-0.036, 0.016] | -0.010 | [-0.054, 0.034] |
|  | Hippocampus | -0.019 | [-0.047, 0.012] | -0.004 | [-0.037, 0.028] |
|  | Dorsal attention network | -0.008 | [-0.024, 0.017] | -0.035 | [-0.076, 0.005] |
|  | Ventral attention network | -0.033 | [-0.064, -0.005] | 0.019 | [-0.023, 0.074] |
|  | Frontoparietal network | -0.013 | [-0.039, 0.019] | -0.036 | [-0.07, -0.003] |
| E-PHATE | Amygdala | 0.091 | [0.062, 0.118] | 0.062 | [0.019, 0.11] |
|  | Hippocampus | 0.110 | [0.079, 0.139] | 0.029 | [-0.017, 0.076] |
|  | Dorsal attention network | 0.076 | [0.052, 0.107] | 0.014 | [-0.027, 0.067] |
|  | Ventral attention network | 0.100 | [0.074, 0.125] | 0.054 | [0.014, 0.097] |
|  | Frontoparietal network | 0.091 | [0.062, 0.124] | 0.047 | [0.004, 0.092] |

### E-PHATE strengthened cross-sectional associations with emotional and behavioral problems

We tested whether E-PHATE strengthened associations between brain function and CBCL scores(30) by comparing the partial Spearman’s p of multiple linear regression models trained on voxel-wise data, PHATE, and E-PHATE embeddings. Voxel-wise data from the 2-back vs. 0-back working memory contrast showed a small effect yet significant relationship to CBCL total problems only in the hippocampus (p = 0.032, 95% CI=[0.005, 0.063]); Table 2). PHATE embeddings of the 2-back vs. 0-back contrast were significantly related to total problems in all ROIs with small-to-moderate effect sizes, but only outperformed voxel data significantly in the frontoparietal network. E-PHATE reflected stronger associations between brain activation and total problems relative to both the voxel data and PHATE embeddings for the 2-back vs. 0-back contrast for every region (corrected p’s all < 0.01, except frontoparietal E-PHATE vs. PHATE p < 0.05). Magnitude of associations between E-PHATE embeddings and overall emotional and behavioral problems were similarly moderate-to-high (p = 0.143 – 0.152) across all regions (Figure 3A; Table 2; Supplemental Data 2).

**Figure 3:**
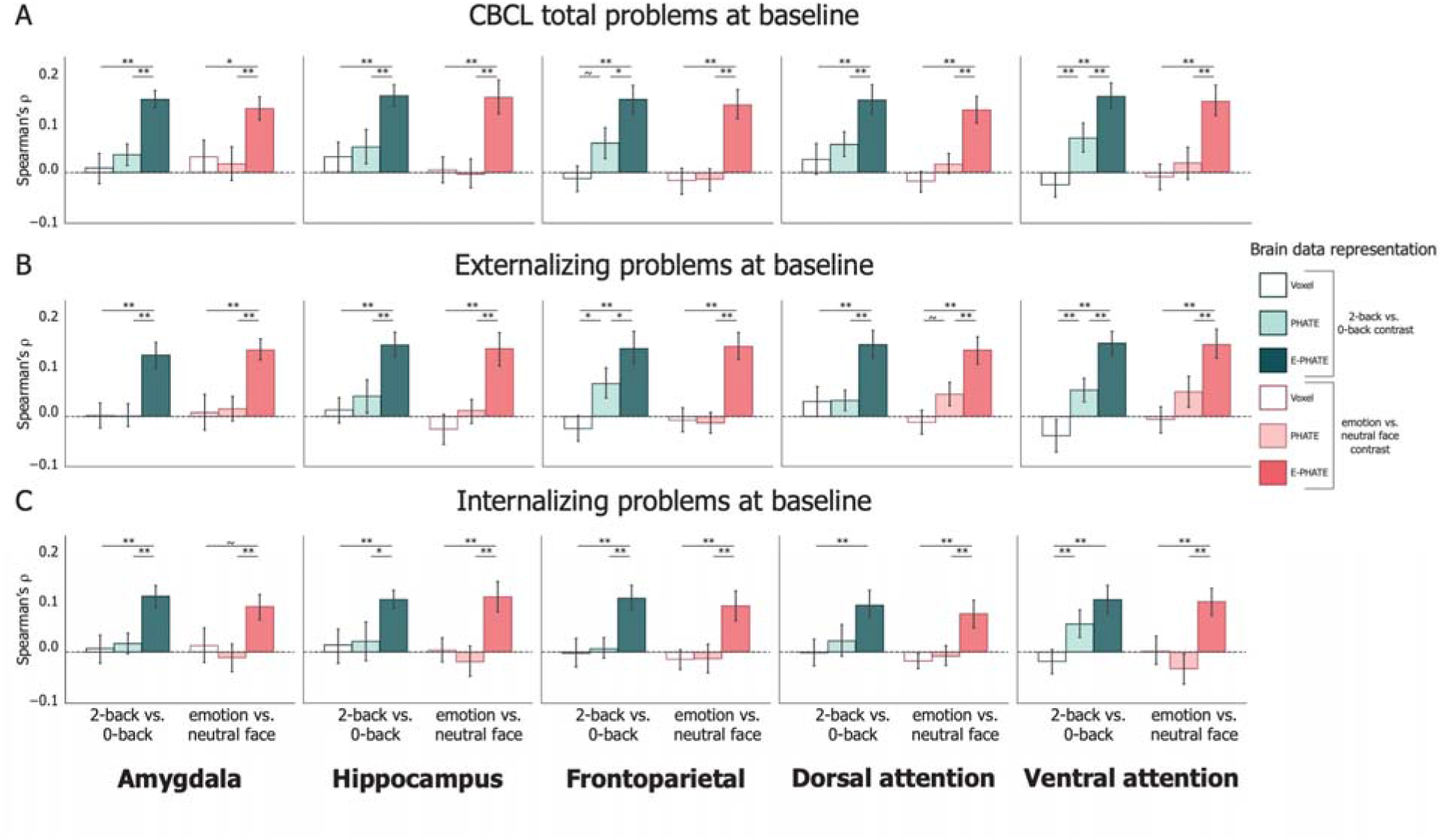
Cross-sectional associations of brain data with mental health problems. Multiple linear regression was used to measure the association between brain data representations and CBCL problem scores and scored as the partial Spearman correlation (p) between true and regression-predicted CBCL scores for held-out participants’ data (20 cross-validation folds). Bars represent the average p across the 20-cross validation test folds. Pairwise permutation tests were used to compute p-values for the difference in p for each cross-validation fold across representations of brain data (i.e., voxel, PHATE, E-PHATE). Both CBCL scores and brain/environmental data were collected at the baseline timepoint. Left bar set in each graph represents the 2-back vs. 0-back contrast; Right bar set in each graph represents the emotion vs. neutral contrast. A=Total problems; B=Externalizing problems; C=Internalizing problems. Error bars represent the 95% confidence interval of the mean p across 20 cross-validation folds. ∼ p < 0.1, * p < 0.05, ** p < 0.01, *** p < 0.001

Replicating previous research showing null associations between emotion-processing activation and emotional and behavioral problems(42), voxel activity from the emotion vs. neutral contrast did not significantly relate to individual differences in CBCL total problems in any ROI. PHATE embeddings of the emotion vs. neutral contrast performed similarly to the voxel data in the strength of its relationship with total problems (all p < 0.035 for voxel and PHATE; small effect size). E-PHATE significantly strengthened this relationship relative to the voxel data and PHATE embeddings in the emotion vs. neutral contrasts for all regions (corrected p’s all < 0.01, except amygdala E-PHATE vs. voxel p < 0.05). The magnitude of E-PHATE’s performance with emotion vs. neutral contrast was comparably strong to E-PHATE’s performance with the 2-back vs. 0-back contrast, with moderate-to-large effect sizes (all p > 0.12) (Figure 3A; Table 2; Supplemental Data 2).

CBCL total problems can be broken into two main broad-band scales: externalizing and internalizing problems. To examine whether the significant relationships between E-PHATE embeddings and total problems were driven by associations with externalizing or internalizing problems specifically, we repeated the analyses above to predict t-scores for each scale independently. E-PHATE embeddings were more strongly related to externalizing problems relative to the voxel data or PHATE embeddings across both task contrasts in all regions (pairwise comparison between E-PHATE and PHATE and E-PHATE and voxel for each region; corrected p’s all < 0.01, except frontoparietal E-PHATE vs. PHATE p < 0.05; Figure 3B; Table 2). As with the total problem score, E-PHATE’s relationship to externalizing problems showed moderate-to-large effect sizes across regions and contrasts. The magnitude of associations with internalizing problems (p = 0.076 – 0.111 across regions and contrasts) was moderate and lower than the larger effect sizes with externalizing problems, yet still significant (95% CI does not contain 0; Table 2). This was not the case for voxel-wise or most of the PHATE embeddings, aside from the PHATE embedding of 2-back vs. 0-back activation in the ventral attention network, which showed a small but significant effect (p =0.056, 95% CI=[0.034, 0.089]) (Figure 3C). Supplemental analyses (Figure S2; Supplemental Data 1) replicated this pattern across the externalizing and internalizing subscale symptoms: E-PHATE embeddings were significantly associated with all subscales with medium-to-large effect sizes, with the strength of the association depending upon the subscale, fMRI contrast, and ROI.

Benchmarking and robustness analyses showed that: 1) in comparing E-PHATE to other dimensionality reduction methods, E-PHATE embeddings significantly out-perform PCA(43,44) and UMAP(45,46) (Supplemental Methods S7 and Figure S3); 2) the increased sensitivity of E-PHATE was attributable to added information specifically about the environment, versus an increase in the quantity of data about each participant (“E-PHATE control”) (Supplemental Methods S8 and Figure S4); 3) the increased sensitivity of E-PHATE was attributable to the nonlinear combination of brain and environment (“PHATE + environment”); and 4) E-PHATE matrices built solely upon neighborhood disadvantage (ADI) or family conflict(47,48) improved associations relative to no environmental information, yet neither afforded as great of an improvement as the five-feature environment view (Supplemental Methods S9 and Figure S4).

### E-PHATE improved longitudinal prediction of emotional and behavioral problems in the frontoparietal network

To evaluate whether the signals highlighted by E-PHATE could enhance our ability to detect brain activation relevant for future emotional and behavioral problems, we asked whether the same embedding of brain and environmental factors at baseline (ages 9–10) could predict emotional and behavioral problems two years later (at ages 11–12). Using a subset of the original participants (N = 2,371 with complete 2-year data), we embedded baseline voxel-wise brain data and environment measures with E-PHATE. Then, we trained multiple linear regressions to predict CBCL total, externalizing, and internalizing problems at the two-year follow-up (controlling for the corresponding baseline CBCL score and scanner serial number during test).

Prediction of CBCL total problems were only significant (though a small effect size) from E-PHATE embeddings of ventral attention network emotion vs. neutral face activation (p=0.044, 95% CI=[0.011, 0.085]); these problems were not predicted by either voxel or PHATE (Figure 4; Table 2; Supplemental Data 1). Externalizing problems were significantly predicted by E-PHATE embeddings of amygdala activation in the 2-back vs. 0-back contrast (moderate effect size; p=0.060, 95% CI=[0.023, 0.096]), and by E-PHATE embeddings of hippocampus and ventral attention network emotion vs. neutral face activation (moderate effect sizes; hippocampus: p=0.053, 95% CI=[0.006, 0.096], ventral attention network: p=0.053, 95% CI=[0.012, 0.090]). Longitudinal internalizing problems were more strongly predicted than externalizing or total problems, with small-to-moderate effects, again only using E-PHATE embeddings (significant in the amygdala, frontoparietal, and attention networks for the emotion vs. neutral face contrast, and the hippocampus, frontoparietal, and dorsal attention networks for the 2-back vs. 0-back contrast; Figure 4A; Supplemental Data 2). Longitudinal predictions were more focused to specific contrasts and regions than cross-sectional associations, possibly suggesting more nuanced, pointed mechanisms related to distinct problems over time.

**Figure 4:**
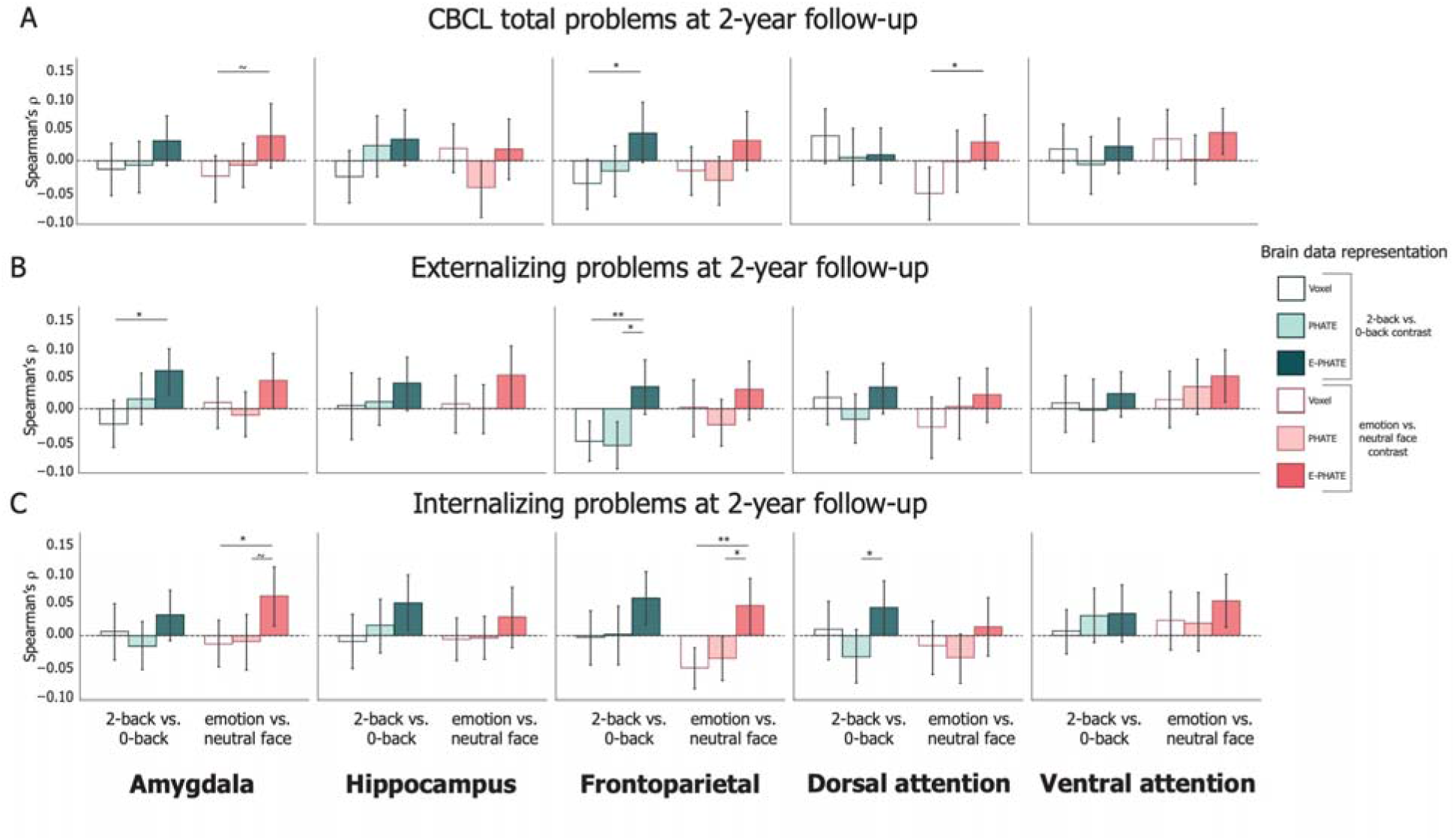
Longitudinal prediction of emotional and behavioral problems. Multiple linear regression was used to predict CBCL problem scores at the 2-year timepoint from baseline brain/environmental data representations and scored as the partial Spearman correlation (p) between true and regression-predicted CBCL scores for held-out participants’ data (20 cross-validation folds). Bars represent the average p across test folds. Pairwise permutation tests were used to compute p-values for the difference in p across representations of brain data (i.e., voxel, PHATE, E-PHATE). Left bar set in each graph represents the 2-back vs. 0-back contrast; Right bar set in each graph represents the emotion vs. neutral contrast. A=Total problems; B=Externalizing problems; C=Internalizing problems. Error bars represent the 95% confidence interval of the mean p across 20 cross-validation folds. ∼ p < 0.1, * p < 0.05, ** p < 0.01, *** p < 0.001

## Discussion

Decades of theories and empirical research indicate that adolescent neurobiology and environmental context interact to shape development of emotional and behavioral problems. Yet, prior work has struggled to capture the complexity of this interplay. Here, we used nonlinear manifold learning to test: 1) whether we can improve the basic associations between adolescent neurobiology and individual differences in cognitive/mental health(21,49), and 2) model brain-environment interactions in a way that reflects their nonlinear multi-dimensionality and relates to mental health outcomes. Across a large, sociodemographic diverse sample of US adolescents, PHATE embeddings enhanced the association of fMRI task activation in multiple brain regions and networks with individual differences in cognitive processing. However, the standard PHATE embeddings did not greatly improve associations with emotional and behavioral problems. Using E-PHATE, we vastly improved detection and prediction of emotional and behavioral problems. Overall, our results demonstrate that manifold learning techniques are well-suited for the complexity of multimodal developmental data and have great potential to enhance research on the neurobiology of emotional and behavioral problems in adolescents.

A major goal of developmental science is to characterize the interplay between adolescents and their broader environments in order to identify early markers of risk and novel targets for intervention(3,4,14,15,28,29). This work offers a substantial methodological advance toward that goal through the development of E-PHATE. E-PHATE offers researchers a data-driven method for capturing the nonlinear interactions of biological—environmental factors, in contrast with prior univariate approaches which have modeled these interactions as a simple product of two variables(50) or multivariate approaches which have been limited in combining brain and environment and focus more on optimizing the brain data for prediction(18,19). By incorporating exogenous information about adolescents’ neighborhoods and families as essential data lending structure to the brain activation manifold, E-PHATE improved associations between brain function and emotional and behavioral problems. Previous studies have questioned the reliability of developmental, task-based fMRI(49) and of empirical support linking specific ROIs (e.g., amygdala) to emotional and behavioral problems in youth(51–53), yet E-PHATE highlighted signals relevant for understanding emotional and behavioral problems in every ROI and network we examined across both contrasts. These results demonstrate that efforts to elucidate relationships between adolescent brain function and emotional and behavioral problems may be stifled if researchers fail to consider the broader context in which the brain develops(29,54,55).

Beyond the applications in developmental neuroscience and clinical psychology outlined above, E-PHATE shows promise for a variety of big-data challenges. E-PHATE addresses a key limitation of many manifold learning methods (e.g., PCA, UMAP, or PHATE), which identify latent structure in a purely unsupervised fashion. In other words, the algorithms do not integrate or evaluate the interplay of multiple measurement types into one latent structure. Other methods (e.g., partial least squares regression, PCA ridge regression, or canonical correlation analysis) used to study brain-behavior associations from high-dimensional data do so by refining latent components with the direct goal of maximizing prediction of a certain variable (e.g., CBCL scores), and then testing those components on out-of-sample data(17–19). In contrast, E-PHATE combines different measurements of the same samples into the manifold calculation but does not refine its representation with any specific goal of downstream prediction; thus, it, maintains both the benefits of unsupervised manifold geometry discovery and hypothesized structure.

The present work should be viewed in light of a few limitations. First, we focused on specific regions and networks that have previously been related to memory- and emotion-processing and mental health(5,6,14,27,35,40,41). Yet, investigating other brain areas or whole-brain approaches may be relevant for understanding task performance and emotional and behavioral problems. Second, manifold learning algorithms are not able to discern the direction or specific patterns of brain activation that contribute to associations with task performance and emotional and behavioral problems. Nonlinear manifolds are also challenging to faithfully extend to untrained samples(56–58), which is an important future direction to increase the impact of this work. Third, while E-PHATE could predict emotional and behavioral problems two years later, results from the current study are correlational and cannot speak to causation. Considering research showing bidirectional relationships between the environment and emotional and behavioral problems, future studies should incorporate manifold learning within other longitudinal designs. Fourth, the results in the current paper only reflect a snapshot of development. Since the peak onset of emotional and behavioral problems is later in adolescence(1), future research investigating a larger developmental window is needed.

In all, we present evidence for a complex interplay between environments, brain function, and emotional and behavioral problems as uncovered by a new, general-purpose method with interdisciplinary applications. Ultimately, data-driven, interdisciplinary approaches that characterize adolescents’ changing neurobiology within the context of their environments may allow us to identify early markers of risk and novel targets for intervention.

## Acknowledgements

We thank Smita Krishnaswamy, Ph.D., for helpful conversation.

This work was supported by NSF GRFP Award 2139841 (E.L.B.) and NIH Grant R21DA057592 (A.B.S.). Data used in the preparation of this article were obtained from the Adolescent Brain Cognitive Development (ABCD) Study (https://abcdstudy.org), held in the NIMH Data Archive (NDA). This is a multisite, longitudinal study designed to recruit more than 10,000 children age 9-10 and follow them over 10 years into early adulthood. The ABCD Study is supported by the National Institutes of Health and additional federal partners under award numbers U01DA041048, U01DA050989, U01DA051016, U01DA041022, U01DA051018, U01DA051037, U01DA050987, U01DA041174, U01DA041106, U01DA041117, U01DA041028, U01DA041134, U01DA050988, U01DA051039, U01DA041156, U01DA041025, U01DA041120, U01DA051038, U01DA041148, U01DA041093, U01DA041089, U24DA041123, U24DA041147. A full list of supporters is available at https://abcdstudy.org/federal-partners.html. A listing of participating sites and a complete listing of the study investigators can be found at https://abcdstudy.org/consortium_members/. ABCD consortium investigators designed and implemented the study and/or provided data but did not necessarily participate in analysis or writing of this report. This manuscript reflects the views of the authors and may not reflect the opinions or views of the NIH or ABCD consortium investigators.

The ABCD data repository grows and changes over time. The ABCD data used in this report came from DOI: <u>10.15154/1523041</u>.

E-PHATE software and all analysis code will be released publicly upon manuscript acceptance.

## Disclosures

The authors report no potential conflicts of interest. This article has been posted as a preprint to bioRxiv.

## Notes

### Competing Interest Statement

The authors have declared no competing interest.

### Summary of Updates

This version has updated longitudinal analyses, expanded benchmarking, and clarified explanations of E-PHATE and cross-validation

