## Supplemental information for "Manifold learning uncovers nonlinear interactions between the adolescent brain and environment that predict emotional and behavioral problems"

**Supplemental materials**

**Supplemental methods**

1. *ABCD Study Participants*

The ABCD Study is a longitudinal study following youth from 21 sites across the United States(1). Study procedures received centralized University of California, San Diego and site-specific institutional review board approval(2). Parents or caregivers provided written informed consent and adolescents gave written assent for study participation. ABCD study-wide exclusion criteria included a diagnosis of schizophrenia, moderate to severe autism spectrum disorder, intellectual disability, substance use disorder at recruitment, persistent major neurological disorders, multiple sclerosis, sickle cell disease, seizure disorders like Lennox-Gastaut Syndrome, Dravet Syndrome, and Landau Kleffner Syndrome.

To determine the current sample, we excluded participants who were lacking: (1) inclusion-recommended EN-back task; (2) environment measures (see below for specific measures); or (3) Child Behavior Checklist (CBCL) scores. We further excluded participants who had missing or extreme beta weights in their task fMRI contrasts (see Section 4 below for more information). Finally, we selected at random one participant per family to be included. To be included in the longitudinal analysis, participants needed to have the environment, CBCL, and EN-back task scores at the 2-year follow-up visit (Figure S1). Participant demographics are included in the main text, Table 1.

We used a logistic regression understand how the selected sample differed from the full ABCD study cohort in terms of the behavioral and environment measures we studies. Given that our research question was centered around the EN-back task activation, we only compared our selected cohort (N=4,732 participants at baseline) to all the participants who had viable EN-back fMRI data (N=7,932; not included N=3,197). The logistic regression was trained to predict whether a participant was included in the cohort or not from their scores on the following: CBCL total problem and externalizing and internalizing subscales; family conflict, family support and acceptance, neighborhood threats (both adult and child rated), and neighborhood deprivation; parental education; and household income (see Methods S2 for more information on these variables
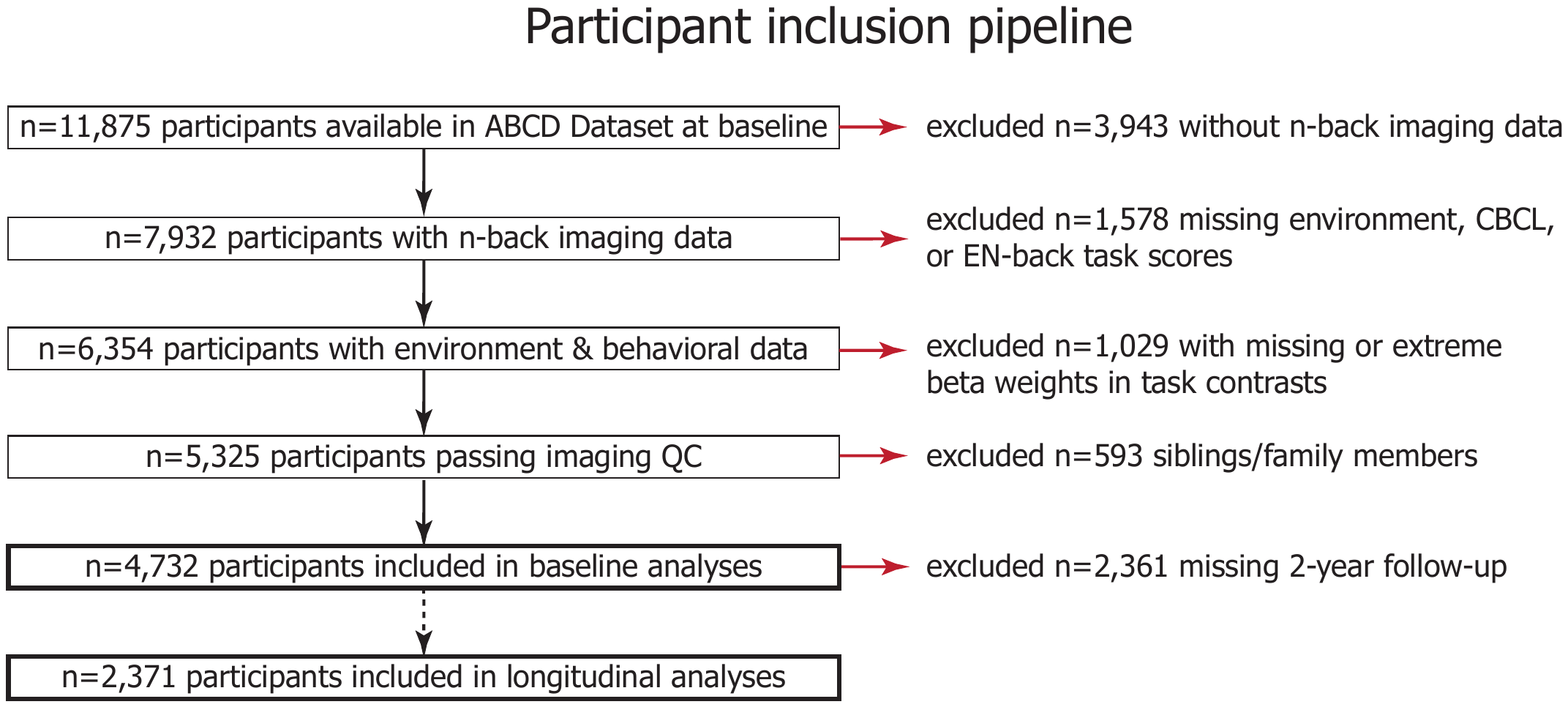
*Figure S1: Flowchart illustrating participant inclusion/exclusion criteria*

2. *Environment measures*

Measures of the environment were selected to characterize youth’s family and neighborhood environments using baseline assessments from the ABCD Study Culture and Environment(3) and Linked External Data(4) protocols.

Family threats were measured with the youth-report ABCD Family Environment Scale–Family Conflict Subscale, which included nine items measuring youth’s perception of anger and conflict expressed among family members. For each item, youth endorsed whether a statement was true or false for most family members. All items were summed with higher scores indicating *more* family conflict.

Family support and acceptance was assessed with the Children’s Report of Parental Behavioral Inventory, which included five items measuring youth’s perception of caregiver acceptance and support. For each item, youth indicated whether a statement was not, somewhat, or a lot like their primary caregiver. All items were averaged and reverse scored such that higher scores indicated *less* family support/acceptance.

Neighborhood threats were measured using the ABCD Neighborhood Safety/Crime Survey, which assessed perceived safety/crime in participants’ neighborhoods. Because the adolescent assessment included only a single item asking youth whether they strongly agreed or strongly disagreed with a statement about their safety from neighborhood crime, neighborhood threats were measured as the mean of all adolescent- and caregiver-report items. The caregiver survey included the exact item from the youth survey with two other items asking whether caregivers strongly agreed or strongly disagreed with statements about their perceived safety while walking and violence being a problem in their neighborhood. All adolescent- and caregiver-report items were reverse-scored such that higher scores indicated *more* neighborhood threats.

Neighborhood deprivation was measured using the Area Deprivation Index (ADI), which is a composite index of neighborhood socioeconomic disadvantage based on income, education, employment and housing quality in a census tract derived from the American Community Survey. Participants’ ADI national percentile were calculated for their primary address, and higher scores indicated *more* neighborhood deprivation.

3. *Emotional n-back task*

Participants performed the emotional n-back (EN-back) task while in the fMRI scanner. The task included two ~5-minute fMRI runs, each with eight task blocks and four 15s fixation blocks. During each run, participants performed four 0-back (low memory load) and four 2-back (high memory load) blocks with happy, fearful, or neutral face images or place images. Stimuli were presented for 2s and followed by a fixation cross. For each trial, stimuli were presented for 2s and followed by 500ms fixation cross. On 0-back blocks, participants were instructed to press “match” when the stimulus was the same as a target stimulus presented at the beginning of the block, and “no match” if not. On 2-back blocks, participants were instructed to press “match” when the stimulus was the same as the stimulus presented two trials back, and “no match” if not.

4. *Neuroimaging data*

fMRI data were collected on Siemens Prisma, Phillips, and GE 750 3T scanners using a 32-channel head coil(1). Functional images were collected with a multiband gradient echo-planar imaging sequence and the following parameters: TR = 800 ms, TE = 30 ms, flip angle = 52°, 60 slices acquired in the axial plane, voxel size = 2.4mm^3^, multiband slice acceleration factor = 6. All fMRI data were preprocessed by the ABCD Study Data Analysis, Informatics, and Resource Center (DAIRC) as detailed elsewhere(5). Participants were excluded based on low quality structural scans, or fewer than 550 degrees of freedom in preprocessed, concatenated timeseries data.

EN-back task activation was estimated for each participant using general linear models with fixation, 2-back and 0-back condition, and happy, fearful, and neutral face and place stimuli as predictorsEmotion processing activation was measured as the linear contrast of emotional (happy and fearful) and neutral face blocks, and cognitive processing activation was measured as the linear contrast of 2-back and 0-back blocks, as in prior studies(6,7). After processing, we excluded contrast maps for: missing over 260 grayordinate values (e.g., missing values outside the medial wall), and extreme values (> 3 SD from the group mean) for the mean or standard deviation of beta weights across all grayordinates, as in prior studies(6).

5. *Exogenous PHATE (E-PHATE) procedure*

Given a matrix $X=x_{1},x_{2},x_{3}\ldots x_{n}$ where$x_{n}\in R^{v}$ for participant *n* is a *V*-dimensional vector, and *V* is the number of voxels or vertices in a ROI. The construction of the PHATE diffusion geometry is summarized as steps 1–4 here and described in more detail elsewhere(8,9):

1. Calculate Euclidean distance matrix between data pairs (pairs of participants), where:

$$D\left( i,j \right)=\left| \left| x_{i}-x_{j} \right| \right|^{2}$$

2. Convert *D* from a distance matrix into a local affinity matrix *K* using an adaptive bandwidth Gaussian kernel. This captures local neighborhoods in the data.

$$\mathcal{K}\left( x,y \right)=\frac{\mathcal{G}\left( x,y \right)}{|\mathcal{G}\left( x,\cdot\right)|_{1}^{\alpha}|\mathcal{G}\left( y,\cdot\right)|_{1}^{\alpha}},\quad\mathcal{G}\left( x,y \right)=e^{-\frac{|x-y|^{\alpha}}{\sigma}}$$

3. Row-normalize *K* to define transition probabilities into the *N* × *N* row stochastic matrix, *P.*

$$P\left( i,j \right)=\frac{\mathcal{K}\left( x_{i},x_{j} \right)}{|\mathcal{K}\left( x,\cdot\right)|_{1}}$$

4. Use probabilities *P* for a Markovian random-walk diffusion process and compute the diffusion timescale *t_D_*, which specifies the number of steps taken in the random walk. *t_D_* is calculated based on the spectral or von Neumann entropy of the diffusion operator. Perform the *t_D_*-step random walk over *P* by raising *P* to the *t_D_* power, resulting in $P^{t_{D}}$. Based on the representation of $P^{t_{D}}$, the diffusion potential distance *P_D_* is calculated between the distributions at the *i*-th and *j*-th rows ($P_{i}^{t_{D}}, P_{j}^{t_{D}})$ as follows:

$$P_{D}\left( i,j \right)=\sqrt{\sum_{k} \left( {\log\left( P^{t_{D}}\left( i,k \right)-P^{t_{D}}\left( j,k \right) \right)}^{2} \right\}}$$

For more details about the PHATE algorithm, we refer readers to Moon et al., 2019(8).

Given a matrix $F=f_{1},f_{2},f_{3},\ldots f_{n}$ where $f_{n}\in R^{k}$ for participant *n* is a *k*-dimension vector, and *k* is the number of exogenous (matched samples of a different measurement) features one opts to include in the second view. The E-PHATE procedure extends beyond the original PHATE formulation in three more steps:

5. Calculate the affinity matrix *A* as the Euclidean distance between data pairs (pairs of participants) across feature vectors, and normalize the value to be between 0 and 1, where 1 corresponds to maximal similarity:

$$A\left( i,j) \right.=\frac{1}{1+||f_{i}-f_{j}||}$$

6. Convert the affinity matrix *A* into the transition probability matrix *P_A_* by row-normalizing the affinity matrix *A*, as in step 3.

$$P_{A}\left( i,j \right)=\frac{\mathcal{A}\left( f_{i},f_{j} \right)}{|\mathcal{A}\left( f,\cdot\right)|_{1}}$$

7. Combine with the result of step 4 via alternating diffusion:

$$P=P_{D}P_{A}$$

8. Embed with metric multidimensional scaling (m-MDS) into *M* dimensions, where $1\leq M \leq3$ for visualization or higher for downstream analysis.

Step 7, the dual diffusion of *P_D_* and *P_A_*, allows for *P* to represent both the transitional probability between samples in terms of the geometry of brain activation (*P_D_*) and in the similarity among exogenous features (*P_A_*). In other words, the dual-diffusion step allows the E-PHATE diffusion matrix to represent data geometry across the neuroimaging dimensions (voxels or surface vertices, within *P_D_*) and across the dimensions of environmental metrics (*P_A_*). After computing *P*, which is our *N* × *N* E-PHATE diffusion matrix, we embed it into *D* dimensions via multidimensional scaling.

For the present analyses, we retain 20 dimensions in our embeddings, across PHATE, E-PHATE, and all benchmarking methods. Voxel-wise data dimensionality for each region is unchanged after extraction from a given parcellation; see Table S2 for voxel dimensions.

6. *Regression and Spearman correlation analyses*

We used multiple linear regression to measure associations between brain data representations (e.g., voxel-wise, PHATE, E-PHATE) and target behavioral measures (i.e., EN-back task performance or CBCL problem scores). Regression analyses were run in two ways: 1) using brain data from the baseline timepoint to predict behavioral scores at the baseline timepoint (“cross sectional associations”), and 2) using brain data from the baseline timepoint to predict behavioral scores at the 2-year follow-up (“longitudinal prediction”).

Regression analyses were conducted as follows for all embedding methods. Consider a matrix $X=x_{1},x_{2},x_{3}\ldots x_{n}$ where $x_{n}\in R^{v}$for participant *n* is a *V* dimensional vector, and *V* is the number of voxels (or vertices) in a region or network. Matrix *X* is embedded into *D* dimensions using a given embedding technique (e.g., PCA, PHATE, E-PHATE), resulting in a new matrix $X_{e}=x_{e_{1}},x_{e_{2}},x_{e_{3}}\ldots x_{e_{n}}$ where $x_{e_{n}}\in R$ for participant *n* is a *D* dimensional vector. After computing *X_e_*, the regression pipeline is as follows. We defined a 20-fold cross-validation procedure to split the participants into train/test folds. Participants were sorted into cross-validation folds randomly, but these splits were kept constant across all regression analyses. At each fold, we trained a multiple linear regression using ordinary least squares to predict training participants’ behavioral scores from their vectors in the embedding space. We then scored the regression model on the unseen testing participants by asking the model to predict test participants’ behavioral scores from their vectors in the embedding space and computing the partial Spearman’s correlation between their predicted and true behavioral scores, including scanner serial number as a covariate. Concerns regarding non-normality among behavioral scores was addressed with a large sample size and by scoring our model using Spearman’s rank order correlation as opposed to Pearson’s correlation(10). This was repeated for each cross-validation fold. Correlation coefficients (ρ) were then averaged across cross-validation folds, resulting in one average score per region/network, contrast, and embedding method.

As a full example, consider using the PCA data matrix for the baseline EN-back 2-back vs. 0-back contrast in the hippocampus to predict CBCL total problem scores. We began with a matrix of dimensions 4,372 samples (participants) by 1,170 features (voxels). We ran PCA over this matrix to reduce the voxel dimensions and retained 20 components, resulting in an embedded matrix of 4,372 samples by 20 features. We then split the 4,372 participants into 20 cross-validation folds. For each fold, we trained a multiple linear regression on the PCA loadings for 4,156 participants to predict their CBCL total problem scores. We then scored this regression on the remaining 216 participants by predicting their CBCL total problem scores from their PCA loadings, then computing a partial Spearman’s correlation between their true and predicted CBCL total problem scores. We included scanner serial number as a covariate in this partial correlation (as some ABCD sites have multiple scanners).

All analyses were conducted in python version 3.12. Multiple linear regression models were implemented using scikit-learn(11) linear regression (scikit-learn.org/stable/modules/generated/sklearn.linear_model.LinearRegression.html). Cross-validation folds were determined using scikit-learn(11) KFold, using a pre-defined random state to assure the same random samples per fold ([scikit-learn.org/stable/modules/generated/sklearn.model_selection.KFold](https://scikit-learn.org/stable/modules/generated/sklearn.model_selection.KFold)). Model accuracy was computed using pinguoin-stats partial_corr package ([pingouin-stats.org/build/html/generated/pingouin.partial_corr.html](https://pingouin-stats.org/build/html/generated/pingouin.partial_corr.html)) which uses the methods described in Kim (2015)(12) to compute partial correlation coefficients and associated p-values, and has been tested for convergence with the R package “ppcor” (cran.r-project.org/web/packages/ppcor/). We used the PHATE implementation released in version 1.0.11 (<https://phate.readthedocs.io/en/stable/>) downloaded with pip.

*7. Dimensionality reduction benchmarks*

We benchmarked dimensionality reduction with E-PHATE against several other dimensionality reduction and manifold learning approaches. Critically, we only compared E-PHATE against data-driven approaches which *reduce dimensionality based on intrinsic data properties* (e.g., PCA, UMAP, PHATE), as opposed to methods which *refine their dimensionality-reduced components* *to maximize prediction* (e.g., partial least squares regression [PLSR], canonical correlation analysis [CCA], or PC-ridge regression). These methods are increasingly used in brain-wide association studies with powerful effects, but their objectives for learning latent components are inherently distinct from the current work in that their goal is to maximize brain-behavior prediction rather than to uncover a latent, intrinsic geometric structure within multivariate data. For this reason, manifold learning is a more suitable approach to our question.

In our main analyses, we showed associations between E-PHATE embeddings and CBCL total, externalizing, and internalizing problem scores relative to PHATE embeddings and high-dimensional, voxel-wise representations, for given regions/networks and task contrasts (Figure 3). In supplement, we benchmarked E-PHATE with standard PCA and UMAP embeddings. PCA implementation used the scikit-learn implementation ([scikit-learn.org/stable/modules/generated/sklearn.decomposition.PCA.html](https://scikit-learn.org/stable/modules/generated/sklearn.decomposition.PCA.html)) and UMAP used the pip installed, published implementation (umap-learn.readthedocs.io)(13) with the default parameters – local neighborhood size of 15 and Euclidean distance metric. The same input data were given to PCA and UMAP (normalized voxel-wise activation vectors for each participant, in a given brain region/network and contrast).

*8. Alternative versions of (E-)PHATE*

We also designed variants of PHATE and E-PHATE to test the specificity of the environment’s impact. To test whether the nonlinear interplay of environment and brain activation afforded by the dual-diffusion step in E-PHATE yields greater insight into emotional and behavioral problems relative to a linear combination of these features, we compared E-PHATE embeddings with a version of PHATE concatenating the five environment measurements as additional features (i.e., participant *n*’s vector would contain *V* voxels + 5 environment features input to PHATE). We refer to this implementation as “PHATE + Features.”

We also tested whether the dual diffusion simply benefits from additional quantity of data as opposed to the relevance of the additional data by building a second exogenous view with “control” features (i.e., hypothesized to be unrelated to mental health; height, weight, age in months, handedness, and number of siblings). We refer to this implementation as “E-PHATE Control.”

*9. Environmental measures in E-PHATE*

Last, we considered whether specific variables about the environment drove E-PHATE’s improved sensitivity to emotional and behavioral problems. We focused specifically on neighborhood disadvantage (ADI) and family conflict, which are commonly considered in isolation as measures of environmental adversity. We did this by calculating new exogenous views in E-PHATE built solely upon between-subject similarity across these single dimensions.

*10. Statistical testing*

We assessed the statistical significance of the differences in mean 𝜌 for pairs of data types (e.g., associations of E-PHATE with CBCL total problems vs. PHATE with CBCL total problems) using pairwise, flip-sign permutation tests (10,000 iterations). This allowed us to compare the means of different embeddings’ 𝜌 distributions (across cross-validation folds) while controlling for within-subject effects (i.e., that the same participants were used for each cross-validation fold across the distributions being compared). All pairwise p-values were reported as Bonferroni corrected, within contrast type but across regions. We also reported bootstrapped confidence intervals around the mean 𝜌 reported for each embedding type, region, and contrast. Bootstrapping was performed with 1,000 resamples. 95% confidence interval of the mean is displayed. ABCD effect sizes are reported as in (14): small= < 0.05, medium = .05-.15, large= .15-.25.

**Supplemental results**

*1. Comparing cohorts*

We compared the cohort included in our analyses (N=4,372) with all participants who had usable EN-back fMRI data at baseline (N=7,932) to gauge if there were biases in their CBCL or environment measures determining who was excluded from our study. Excluded participants scored significantly lower on the caregiver reported neighborhood safety and crime (β = -0.0567, s.e. = 0.026, z = -2.171, P>|z| = 0.03, CI = [-0.108, -0.006]), indicating that their neighborhoods were safer/had less crime. Excluded participants also scored significantly higher on the area deprivation index (β = 0.0639, s.e. = 0.026, z = 2.485, P>|z| = 0.014, CI = [0.013, 0.115]), indicating their neighborhoods experience greater deprivation. These two scales were the only ones that differed significantly (p < 0.05, uncorrected). Results for all measures in the baseline cohort are in Table S3. When comparing the participants included in the longitudinal cohort (N=2,371 of the original 4,372) with the full EN-back cohort, the cohorts only differ significantly in terms of race/ethnicity. Results for all measures in the baseline cohort are in Table S4.

*2. Associations with CBCL externalizing and internalizing subscales*

We replicated our analyses looking at cross-sectional broad-band scale associations (i.e., CBCL externalizing and internalizing problems) with E-PHATE embeddings for each of the externalizing and internalizing subscale symptoms. Replicating our main result, E-PHATE embeddings were significantly associated with all subscale symptoms, with the strength of the association depending upon the subscale, fMRI contrast, and ROI (all associations showing moderate-to-large effect sizes). As with the broad-band scales, E-PHATE embeddings more strongly reflected externalizing symptoms (aggression, rule-breaking) than they do internalizing symptoms (anxious/depressed, withdrawn/depressed, somatization). These associations were not present in the voxel-wise data for any region or contrast (aside from dorsal attention network 2-back vs. 0-back activation—which showed a small effect—the 95% confidence interval of remaining associations contained 0.

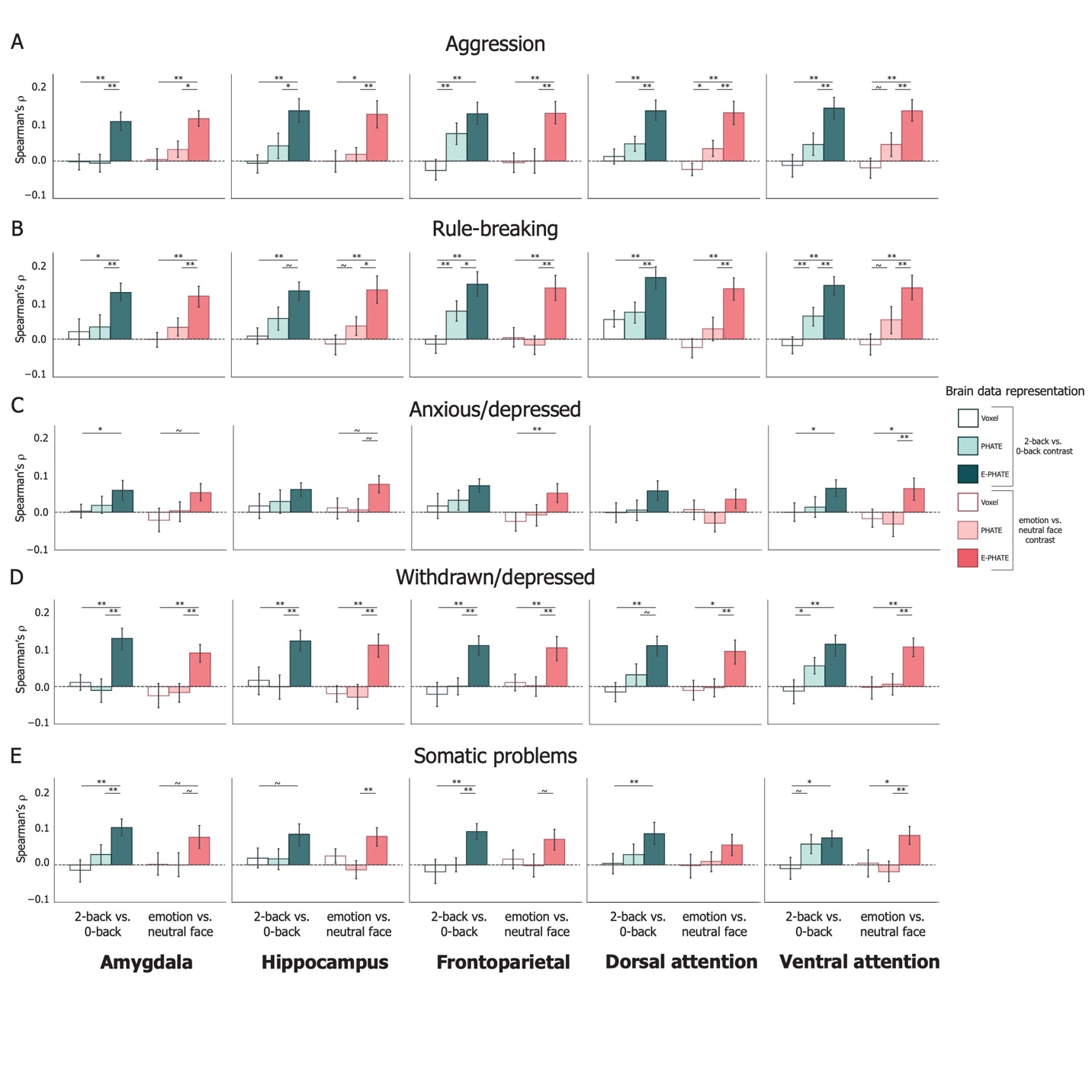

### Figure S2: Association with CBCL subscale scores at baseline timepoint

Aggression and rule-breaking are subscales of the CBCL externalizing problems. Anxious/depressed, withdrawn/depressed, and somatic problems are subscales of the CBCL internalizing problems. Bars represent average partial Spearman’s ρ between model predicted and true behavioral scores on held-out participants’ data, including scanner serial number as a covariate in the scoring. Error bars represent the 95% confidence interval of the mean across 20 cross-validation folds.

~ p < 0.1, * p < 0.05, ** p < 0.01, *** p < 0.001

*3. Benchmarking the associations yielded by E-PHATE*

Results comparing associations of PCA, UMAP, and E-PHATE embeddings with total, externalizing, and internalizing problems across the 5 regions and 2 contrasts (Figure S3). UMAP representations were not significantly associated with emotional and behavioral problems in any region or contrast. PCA yielded weak results comparable to the voxel resolution data included in main analyses (Main Figure 3), with associations with total problems and externalizing problems showing small effect sizes in the frontoparietal and attention networks during the 2-back vs. 0-back contrast.

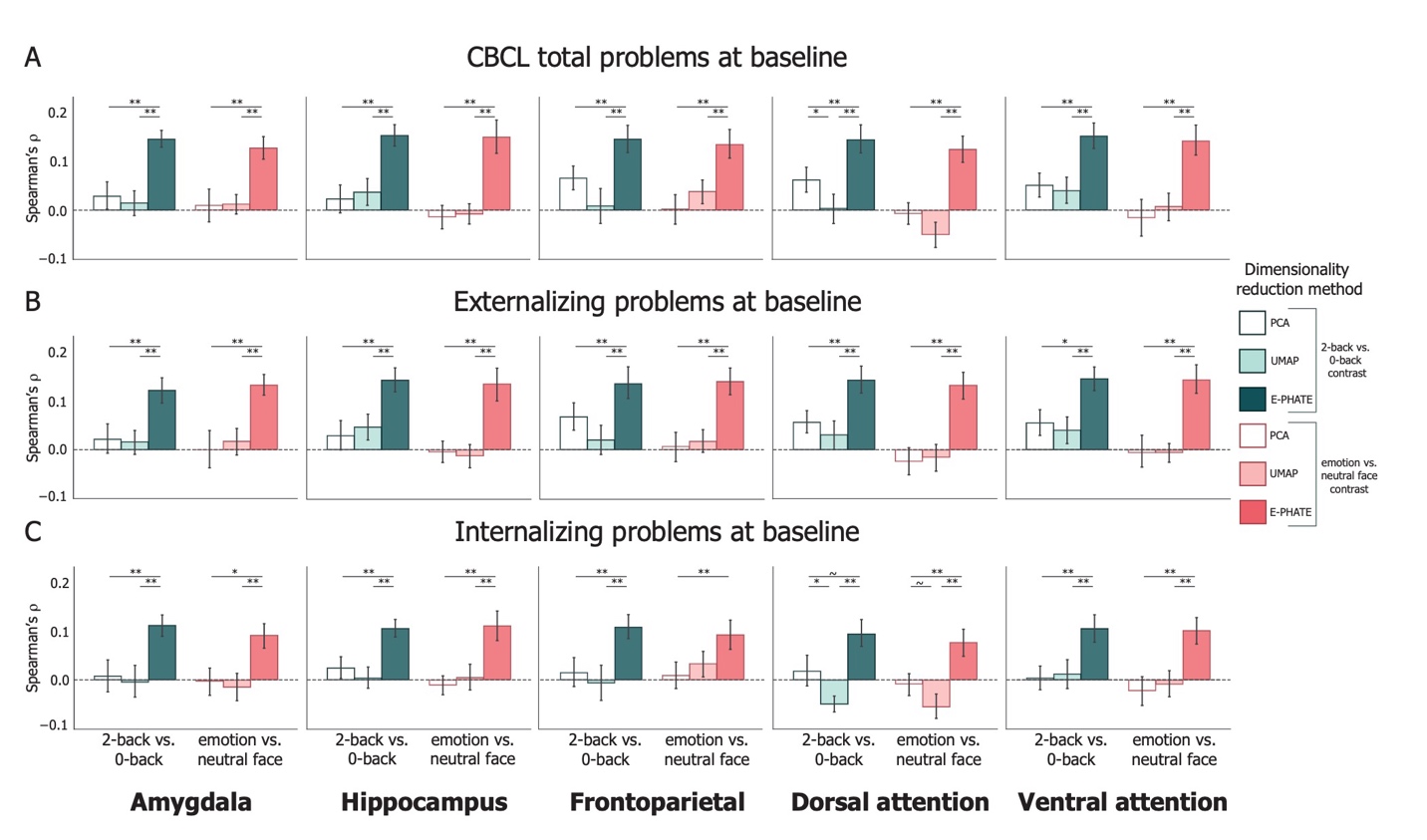
*Figure S3: Common dimensionality reduction methods are less associated with emotional and behavioral problems than E-PHATE embeddings*

Alternative versions of (E-)PHATE tested the specificity of the dual-diffusion approach, which captures nonlinear relationships among the environment and brain in two separate views informing one overall space, with (1) a linear combination of brain and environment information before embedding with PHATE (PHATE + Features) and (2) a control version of E-PHATE, where the exogenous view was built off five variables of a participant’s data hypothesized to be irrelevant to their brain or behavioral data. The linear combination of features (PHATE + Features) matched performance with E-PHATE in the amygdala and the hippocampus (moderate-to-large effect sizes), but across the cortical networks this information weakened associations significantly (zero-to-small effects). It is possible that the environment contributes differently to different regions/networks, as such distinct modeling techniques may or may not be comparable. The irrelevant information included in E-PHATE Control hindered associations between the brain and emotional and behavioral problems, confirming that E-PHATE’s benefit is not driven by a sheer increase in data quantity (Figure S4).

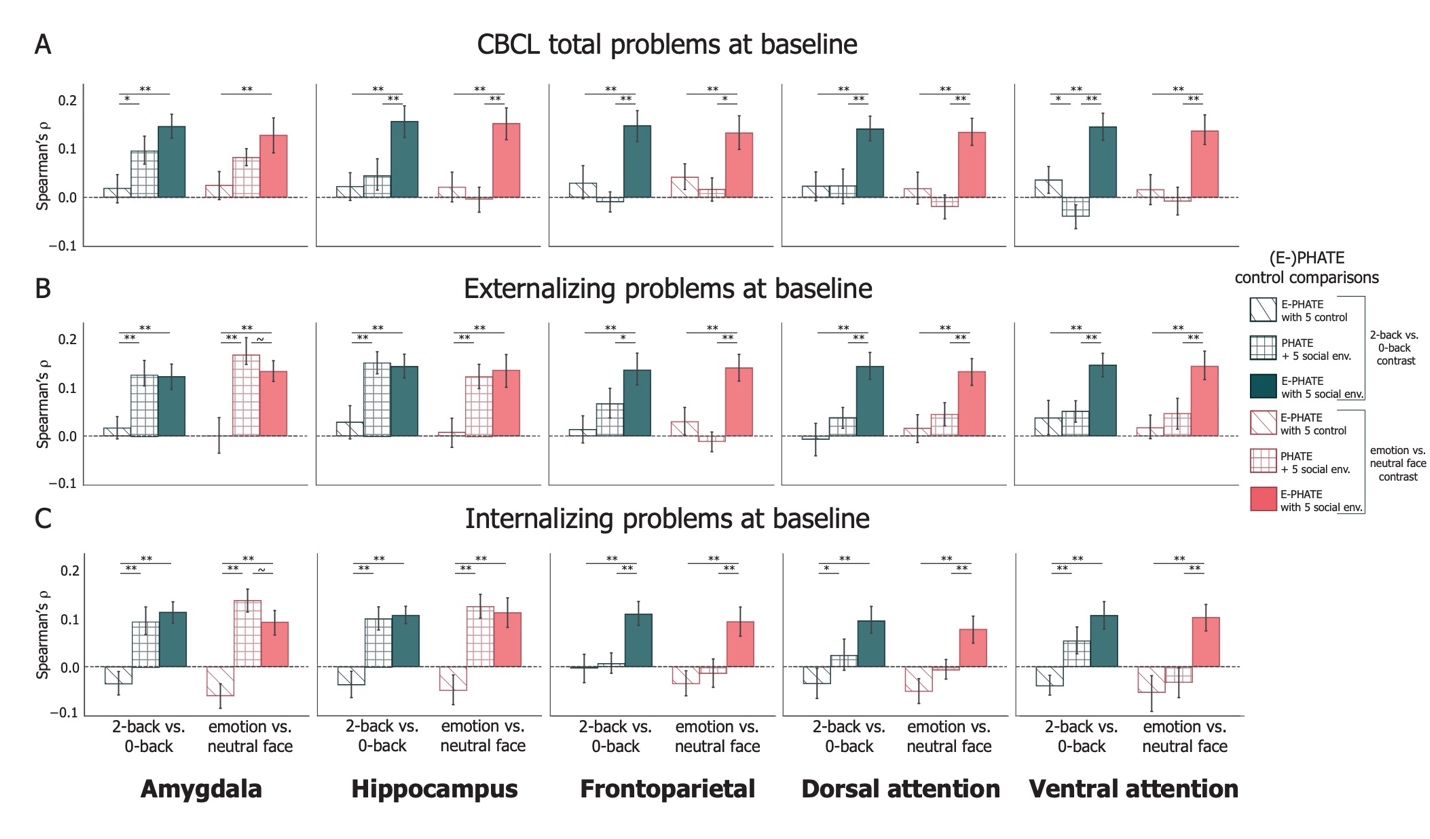

### Figure S4: Altered versions of (E-)PHATE show weaker and less consistent associations with emotional and behavioral problems compared to E-PHATE

Our final benchmark considered the relative contribution of different environment measures in improving E-PHATE’s performance. We tested exogenous factor views of E-PHATE with only neighborhood disadvantage (ADI) and with only family conflict (FES). While adding either neighborhood disadvantage or family conflict alone improved associations with emotional and behavioral problems relative to no environmental information (voxel-wise or PHATE embedding), neither variable in isolation afforded as great an improvement as the five-factor environment view (Figure S5).

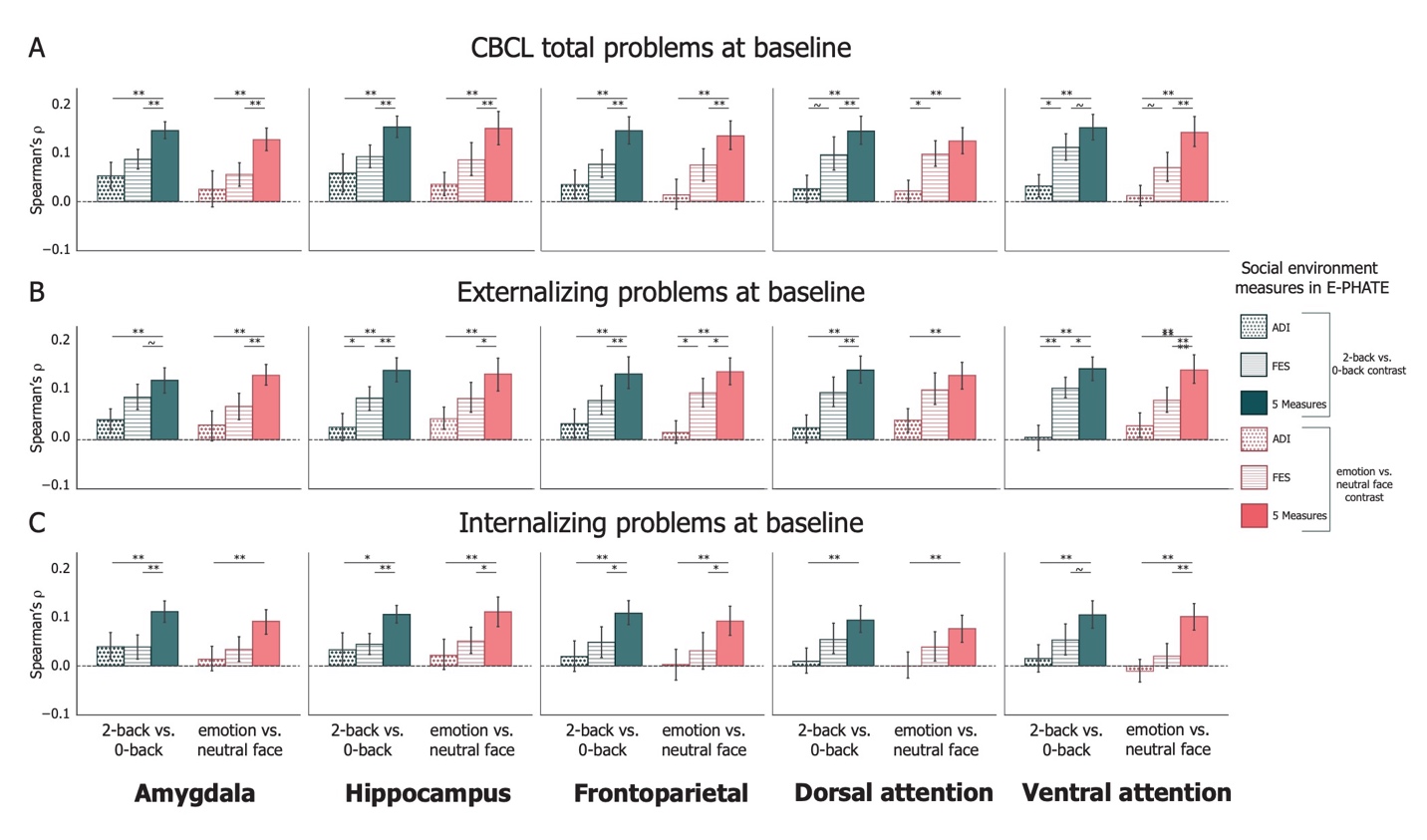

### Figure S5. Neighborhood disadvantage and family conflict individually inform emotional and behavioral associations, but to a lesser degree than all 5 environmental measures.

**Supplemental tables**

*Supplemental Table 1: ABCD Variable Information*

| File name (*.txt) | Variable Name | Description |
| --- | --- | --- |
| abcd_nsc01 | Neighborhood_crime_y | ABCD Youth Neighborhood Safety/Crime Survey |
| abcd_rhds01 | reshist_addr1_adi_perc | Residential history derived - Area Deprivation Index: national percentiles |
| abcd_sscey01 | fes_y_ss_fc | Conflict Subscale from the Family Environment Scale Sum of Youth Report |
|  | crpbi_y_ss_caregiver | Acceptance Subscale Mean of Report by Secondary Caregiver by youth |
| abcd_sscep01 | nsc_p_ss_mean_3_items | Neighborhood Safety Protocol: Mean of Parent Report |
| acspsw03 | race_ethnicity | Child’s race/ethnicity reported by parents |
|  | rel_family_id | Family ID |
| pdem02 | demo_comb_income_v2 | Combined family income |
|  | demo_prnt_ed_v2 | Highest education level of parent |
|  | sex | Participant sex at birth |
| abcd_imgincl01 | imgincl_nback_include | Participant recommended inclusion for EN-back imaging data |
| abcd_mrinback02 | tfmri_nb_all_beh_ctotal_rate | In-scanner EN-back d’ |
| abcd_lt01 | site_id_l | Site ID |
| abcd_mri01 | mri_info_deviceserialnumber | MRI serial number |

*Table S2: Region/network voxel dimensionality*

|  | Region/network name | | | | |
| --- | --- | --- | --- | --- | --- |
|  | *Amygdala* | *Hippocampus* | *Frontoparietal network* | *Dorsal attention network* | *Ventral attention network* |
| **#voxels/vertices** | 573 voxels | 1170 voxels | 4711 vertices | 6427 vertices | 7593 vertices |
| **Atlas citation** | Tian et al., 2020 | Tian et al., 2020 | Yeo et. Al., 2011 | Yeo et al., 2011 | Yeo et al., 2011 |

*Table S3: Logistic regression results, comparing participants included in baseline analyses with all participants with EN-back imaging data.*

|  | **β** | **Std. err.** | **z** | **P > \|z\|** | **[0.025** | **0.975]** |
| --- | --- | --- | --- | --- | --- | --- |
| **cbcl_scr_syn_totprob_t** | -0.0252 | 0.071 | -0.354 | 0.723 | -0.165 | 0.114 |
| **cbcl_scr_syn_external_t** | -0.0231 | 0.048 | -0.480 | 0.631 | -0.118 | 0.071 |
| **cbcl_scr_syn_internal_t** | 0.0600 | 0.048 | 1.250 | 0.211 | -0.034 | 0.154 |
| **reshist_addr1_adi_perc** | 0.0639 | 0.026 | 2.458 | **0.014** | 0.013 | 0.115 |
| **crpbi_y_ss_caregiver** | -0.0231 | 0.025 | -0.919 | 0.358 | -0.073 | 0.026 |
| **fes_y_ss_fc** | 0.0039 | 0.026 | 0.154 | 0.877 | -0.046 | 0.054 |
| **nsc_p_ss_mean_3_items** | -0.0567 | 0.026 | -2.171 | **0.030** | -0.108 | -0.006 |
| **race_ethnicity** | 0.0310 | 0.025 | 1.258 | 0.209 | -0.017 | 0.079 |
| **demo_prnt_ed_v2** | -0.0283 | 0.026 | -1.078 | 0.281 | -0.080 | 0.023 |
| **demo_comb_income_v2** | 0.0409 | 0.025 | 1.655 | 0.098 | -0.008 | 0.089 |

*Table S4: Logistic regression results, comparing participants included in longitudinal analyses with all participants with EN-back imaging data*

|  | **β** | **Std. err.** | **z** | **P > \|z\|** | **[0.025** | **0.975]** |
| --- | --- | --- | --- | --- | --- | --- |
| **cbcl_scr_syn_totprob_t** | 0.0456 | 0.072 | 0.630 | 0.529 | -0.096 | 0.187 |
| **cbcl_scr_syn_external_t** | -0.083 | 0.049 | -1.696 | 0.09 | -0.179 | 0.013 |
| **cbcl_scr_syn_internal_t** | 0.0469 | 0.048 | 0.97 | 0.332 | -0.048 | 0.142 |
| **reshist_addr1_adi_perc** | 0.0435 | 0.026 | 1.658 | 0.097 | -0.008 | 0.095 |
| **crpbi_y_ss_caregiver** | 0.0028 | 0.025 | 0.112 | 0.911 | -0.047 | 0.052 |
| **fes_y_ss_fc** | 0.0267 | 0.026 | 1.046 | 0.295 | -0.023 | 0.077 |
| **nsc_p_ss_mean_3_items** | 0.0041 | 0.026 | 0.155 | 0.877 | -0.048 | 0.056 |
| **race_ethnicity** | -0.0485 | 0.025 | -1.977 | **0.048** | -0.097 | 0.00 |
| **demo_prnt_ed_v2** | -0.0033 | 0.026 | -0.126 | 0.900 | -0.055 | 0.048 |
| **demo_comb_income_v2** | 0.0118 | 0.026 | 0.459 | 0.646 | -0.039 | 0.062 |
